# IKKα kinase coordinates BRD4 and JAK/STAT signaling to subvert DNA damage-based anticancer therapy

**DOI:** 10.1101/2023.06.13.544711

**Authors:** Irene Pecharromán, Laura Solé, Daniel Álvarez-Villanueva, Teresa Lobo-Jarne, Josune Alonso-Marañón, Joan Bertran, Yolanda Guillén, Ángela Montoto, María Martínez-Iniesta, Violeta García-Hernández, Gemma Giménez, Ramon Salazar, Cristina Santos, Marta Garrido, Eva Borràs, Eduard Sabidó, Ester Bonfill-Teixido, Raffaella Iurlaro, Joan Seoane, Alberto Villanueva, Mar Iglesias, Anna Bigas, Lluís Espinosa

**Author notes:** Both first co-authors (Irene Pecharromán, Laura Solé). Mail to.

## Abstract

Activation of the IKK kinase complex has recurrently been linked to colorectal cancer (CRC) initiation and progression. However, identification of downstream effectors other than NF-κB has remained elusive.

Analysis of IKK-dependent substrates after UV-treatment revealed that BRD4 phosphorylation by IKKα is required for chromatin-binding dynamics upon damage. Moreover, IKKα induces the NF-κB-dependent transcription of LIF leading to STAT3 activation, association of BRD4 to STAT3 and recruitment to specific target genes. IKKα abrogation results in defective BRD4 and STAT3 function leading to irreparable DNA damage and apoptotic cell death upon different stimuli. Simultaneous inhibition of BRAF-dependent IKKα activity or BRD4 and the JAK/STAT pathway enhanced the therapeutic potential of 5-FU plus irinotecan in CRC cells, and is curative in a chemotherapy-resistant CRC xenograft model. Coordinated expression of LIF and IKKα is a poor prognosis marker for CRC patients.

Our data uncover a functional link between IKKα, BRD4 and JAK/STAT signaling with clinical relevance.

## INTRODUCTION

Colorectal Cancer (CRC) accounts for about 12% of all deaths from cancer in the European countries (https://ec.europa.eu/eurostat/statistics-explained/index.php/Cancer_statistics_-_specific_cancers#Colorectal_cancer), which highlights the need for innovative therapeutic options. After years of research on cancer and cancer therapy, first line treatment for CRC is still surgery followed by radio- and/or chemotherapy for advance tumors. More recently, and based on the importance of mitogen-activated protein kinase (MAPK) pathway in CRC, antibodies targeting the upstream regulator epidermal growth factor receptor (EGFR) have been included as a second therapeutic option. However, these therapeutic strategies are primarily restricted to the subset of tumors not carrying KRAS or BRAF mutations^1,2^.

Nuclear factor kappa B (NF-κB) signaling has been recurrently identified as a potent tumor driver and the essential linkage between inflammation and cancer. This circumstance led to investigate inhibitors of the Inhibitor of kappa B kinase (IKK) complex, the bottle-neck of NF-κB activation, as potential anticancer agents. However, this possibility was rapidly dismissed due to the extremely high toxicity of general NF-κB inhibition (review in^3^), thus uncovering the need for identifying targetable elements of NF-κB signaling specific of cancer. In this context, the IKKα kinase, which is dispensable for canonical NF-κB signaling, has been the focus of intense investigation (review in^4^). Recently, we demonstrated that IKKα promotes therapy resistance in CRC by facilitating activation of the DNA-damage repair (DDR) pathway through direct phosphorylation of ataxia telangiectasia mutated (ATM) kinase. Inhibition of IKKα activity by vemurafenib or AZ628^5^, or IKKα genetic deletion precluded ATM-dependent DDR following chemo- or radio-therapy treatment leading to specific eradication of cancer cells in vitro and in vivo^5^.

Bromodomain 4 (BRD4) is a member of the bromodomains and extra-terminal (BET) domain family of proteins, which facilitate polymerase II-dependent transcription of genes through recognition of specific histone acetylation marks^6^. BRD4 is unique among the human BET family proteins because it contains a carboxyl-terminal domain (CTD) that interacts with the cyclin T1 and CDK9 subunits of positive transcription elongation factor b (pTEFb) complex to modulate Polymerase II activity^7^. Moreover, BRD4 contains intrinsic Histone Acetyl Transferase activity, which inflicts chromatin de-compaction at its target genes^8^. Additionally, BRD4 facilitates Homologous Recombination and Non-Homologous-End -Joining DNA repair in cancer cells by direct regulation of DDR elements such as CHK1^9–13^ while acting as chromatin insulator to limit the extent of ATM-induced signaling after damage^14^.

BRD4 is functionally involved in cancer development (reviewed in^15^) and was found to impose a proliferative and anti-apoptotic state to mouse embryonic fibroblasts (MEF) and cancer cell lines^16^. BET inhibitors, such as JQ1, have reliably demonstrated their efficacy as anticancer agents and are currently under evaluation in several clinical trials (reviewed in^17^).

JAK/STAT pathway is a well-known tumor driver in different cancer subtypes (reviewed in ^18^), which promotes cell proliferation and therapy resistance by different mechanisms including transcriptional activation of *MYC*^19^, *CYCLIN D1* ^20^, *MDM2* ^21^ and several antiapoptotic genes (review in^22^). JAK/STAT signaling is induced by growth factors and cytokines such as Interleukin 6 (IL6) or the Leukemia Inhibitory Factor (LIF)^23^. JAK-mediated tyrosine (Y) phosphorylation of STAT (Y705 for STAT3) leads to STAT dimerization, nuclear translocation and specific gene transcription^24^. LIF has been described to promote the development and progression of numerous solid tumor types^21,25,26^. It can induce the self-renewal of cancer stem cells/cancer-initiating cells^27^ and promote immune suppression in tumors^28,29^.

We now show that IKKα coordinates BRD4 and STAT3 functions through i) direct phosphorylation of BRD4 thus regulating chromatin-binding dynamics and ii) NF-κB-mediated transcriptional induction of LIF leading to STAT3 activation, formation of BRD4 and STAT3 complexes, and BRD4 recruitment to a subset of STAT3 target genes. Simultaneous inhibition of JAK/STAT and nuclear IKKα activity with BRAF inhibitors^5^ or BRD4 with JQ1 enhanced the anticancer activity of chemotherapy in human CRC cells and tumor xenografts. We identify LIF together with IKKα as a prognosis marker for CRC patients.

Together our data offer new insights on IKKα function in normal and tumor cells and provide novel targets for personalized anticancer therapies.

## RESULTS

### BRD4 S1117 is a phosphorylation substrate of the IKKα kinase

By mass spectrometry (MS) analysis of control and IKKα knocked-down HT29 CRC cells treated with UV^5^, we identified serine (S) 1117 of BRD4 as a UV-inducible and IKKα dependent phosphorylated residue (Figure 1A). This observation raised the possibility that IKKα directly phosphorylates BRD4. By in vitro kinase assay, we confirmed that recombinant active IKKα was able to phosphorylate a BRD4 fragment comprising amino acids 1037 to 1201 (Figure 1B). Mutation of S1117 into alanine precluded BRD4 phosphorylation induced by IKKα (Figure 1C). We identified a second BRD4 residue (S601) whose phosphorylation increased after UV treatment independently of the IKKα status (Figure 1D). Accordingly, recombinant IKKα failed to phosphorylate two different BRD4 fragments containing S601 (Figure 1E).

**Figure 1.**
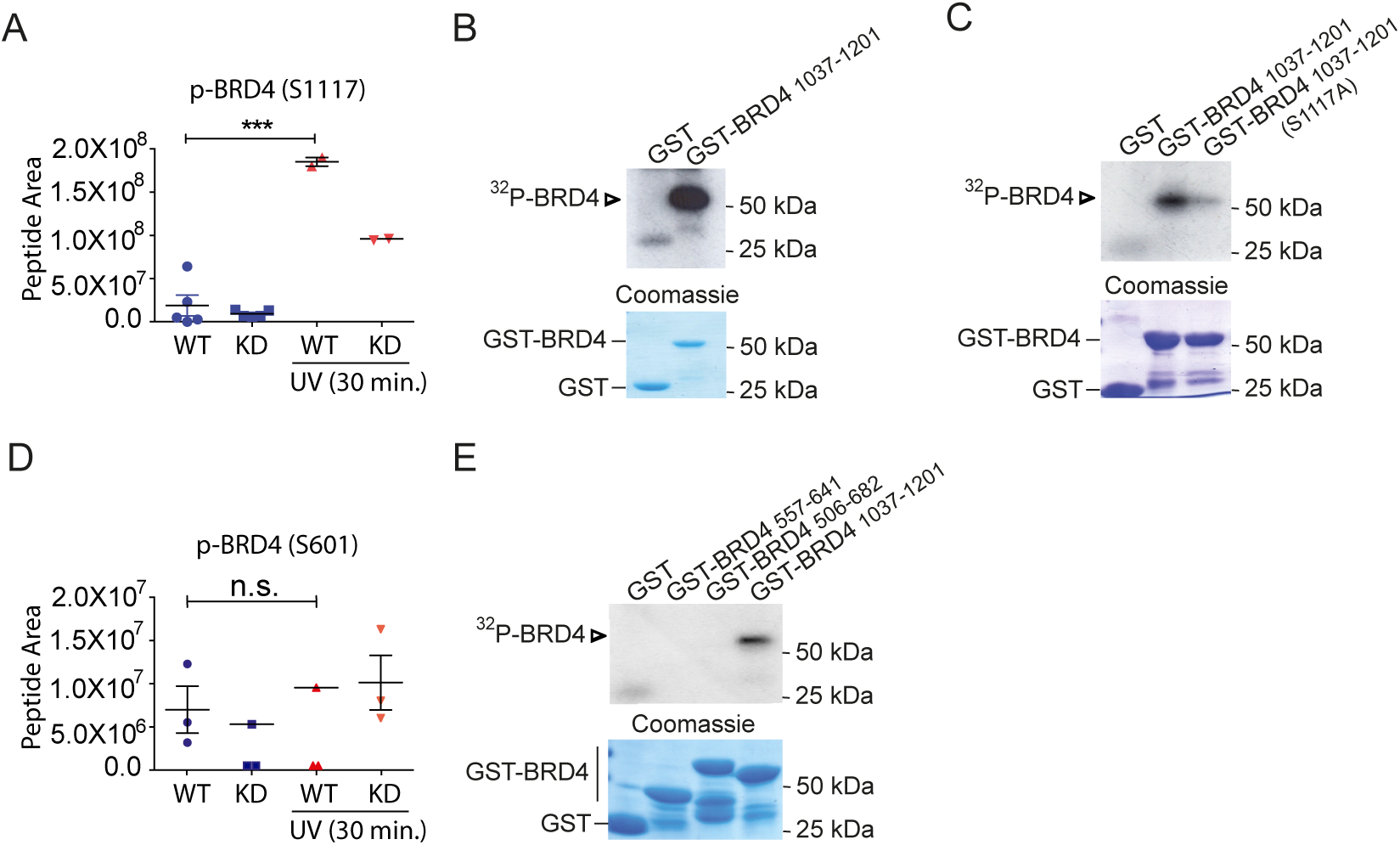
BRD4 S1117 is a phosphorylation substrate of the IKKα kinase. **(A)** Representation of the area corresponding to the phosphor-peptide containing S1117 of IKKα as determined by mass spectrometry analysis (MS) of control and IKKα knocked-down HT29 cells untreated or exposed to UV light (130 mJ) for 30 minutes (n = 5 (untreated) and 2 (treated) biologically independent replicates). **(B, C)** In vitro kinase assay with recombinant IKKα and purified glutathione S-transferase (GST), GST-BRD4 (amino acids [aa] 1037–1201) (B) or the same GST-BRD4 fragment including a S>A mutation at residue 1117 (C) (from one out of three independent experiments). **(D)** Representation of the area corresponding to the phosphor-peptide including S601 from BRD4 as determined by mass-spectrometry (MS) analysis of control and IKKα-knock-down HT29 cells untreated or exposed to UV light (130 mJ) for 30 minutes (n = 3 biologically independent replicates). **(E)** *In vitro* kinase assay with recombinant IKKα and purified GST, two different GST-BRD4 fragments including S601 of BRD4 or GST-BRD4 S1117 [aa1037-1201] as positive control (from one out of three independent experiments).

These results indicate that IKKα specifically phosphorylates BRD4 at S1117. The high conservation of this residue among species (Figure S1) further suggested its functional relevance.

### Two different domains of BRD4 mediate binding to IKKα

By immunoprecipitation (IP) assay of UV-treated cancer cells with the antibody targeting IKKα(p45), we demonstrated the physical binding of IKKα to BRD4 that was slightly increased at 5 minutes of UV exposure and maintained after 15 minutes (Figure 2A). The inducible nature of the IKKα and BRD4 interaction was confirmed by BioID assay using an IKKα protein fused to a biotin donor (see methods). Specifically, upon association with Bio-IKKα, BRD4 become biotinylated and is subsequently recovered in streptavidin precipitates. We observed a significant enrichment in the amount of biotinylated BRD4 after 15 minutes of UV treatment of Bio-IKKα expressing HCT116 cells, which decreased at later time points (i.e. 60 minutes) (Figure 2B). We further confirmed the IKKα and BRD4 interaction by reciprocal IP with the BRD4 antibody (Figure 2C).

**Figure 2.**
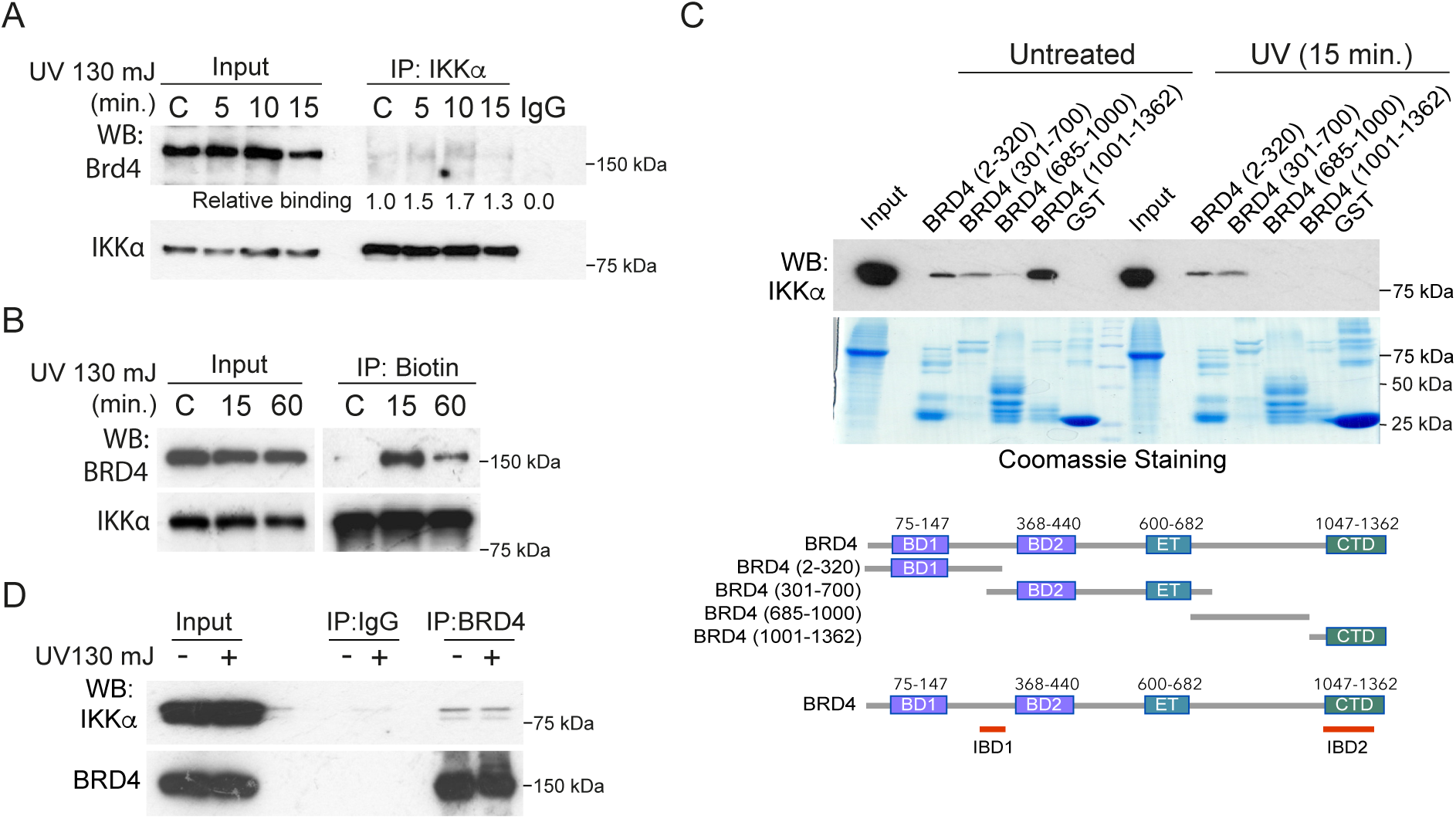
Two different domains of BRD4 mediate IKKα binding. **(A)** Immunoprecipitation assay (IP) with anti-IKKα (p45) antibody from CRC HT29 cells exposed to UV light (130 mJ) and collected at the indicated time points (from one out of three independent experiments). **(B)** Cells lysates from HCT116 cells expressing a fusion BioID-IKKα protein were precipitated using streptavidin-coated beads and then analyzed by western blot with the indicated antibodies (from one out of three independent experiments). **(C)** IP assay with anti-BRD4 antibody from HT29 cell line untreated or collected 15 minutes after UV light exposure (130 mJ) (from one out of three independent experiments). **(D)** Pull-Down assay (PD) of cell lysates from HT29 cell line untreated or collected 15 minutes after UV light exposure (130 mJ) using GST fused to the indicated fragments of BRD4 as bait. Schematic representation of the fragments used for mapping the domains of BRD4 involved in IKKα binding (IKKα Binding <u>D</u>omains 1 and 2) (from one out of three independent experiments). Inputs in A, B and C correspond to 1/25 of the precipitated lysate.

We next mapped the interaction domain of BRD4 to IKKα by pull down assay using various GST/BRD4 fusion protein fragments. Our results indicated that IKKα binds to two different regions of BRD4, the first-one involving amino acids 301 to 320 and the second-one corresponding to the carboxyterminal domain (CTD), which is important for BRD4-mediated transcriptional activation^30^ and includes the IKKα phosphorylation residue 1117 (Figure 2D). Interestingly, activated IKKα (recovered 15 minutes after UV treatment) failed to stably associate to the CTD fragment (Figure 2D) consistent with the transient binding of a kinase with its substrate, which may not necessarily represent the actual binding-dynamics between IKKα and BRD4 in vivo.

### IKKα-induced phosphorylation regulates BRD4 chromatin-binding dynamics and impacts on activation of the DDR elements ATM and Chk1

Association of BRD4 with the chromatin follows a cycling pattern after exposure to DNA damaging agents, being chromatin dissociation and reassociation essential for BRD4-mediated gene transcription^31^. We confirmed the cycling nature of BRD4 chromatin binding upon exposure to different damaging agents including UV light, ionizing radiation (IR) and chemotherapy (5-FU+Iri) in different cell lines (Figures 3A-D and S2A-C). Chromatin dissociation and reassociation of BRD4 display different dynamics in the several models tested and it was severely compromised in IKKα-deficient cells (Figures 3A-B) and in cells treated with the BRAF inhibitor AZ628 (Figure 3C), which prevents IKKα(p45) activation by damage^5^. However, histone acetylation, that is one of the factors regulating BRD4 chromatin binding, was not significantly affected by IKKα deletion or BRAF inhibition (Figures 3A-C and S2D). To test the impact of IKKα-induced BRD4 phosphorylation in chromatin-binding dynamics, we generated knock-in HCT116 cells carrying a BRD4 mutant in which S1117 has been changed to A (BRD4_S1117A_, see methods). BRD4_S1117A_ mutation imposed a massive accumulation of BRD4 in the soluble nuclear compartment and impaired chromatin-binding dynamics when compared with WT BRD4 (Figure 3D). Remarkably, BRD4_S1117A_ HCT116 cells showed defective activation of the DDR elements ATM, ATR and CHK1 upon UV treatment, and increased apoptosis as determined by cleaved caspase 3 levels (Figure 3E), a phenotype that paralleled the effect of full BRD4 inhibition by JQ1 treatment in these cells (Figure S2E). These results suggest that IKKα-mediated phosphorylation of BRD4 at S1117 is required for its correct chromatin binding dynamics and function, including DDR. By qPCR analysis, we found that expression of several DDR-related elements remained unaffected, or even increased, in cells carrying the BRD4_S1117A_ mutant, supporting the concept that BRD4 function on DDR is gene-expression independent^9,32^ but dependent on IKKα-induced phosphorylation.

**Figure 3.**
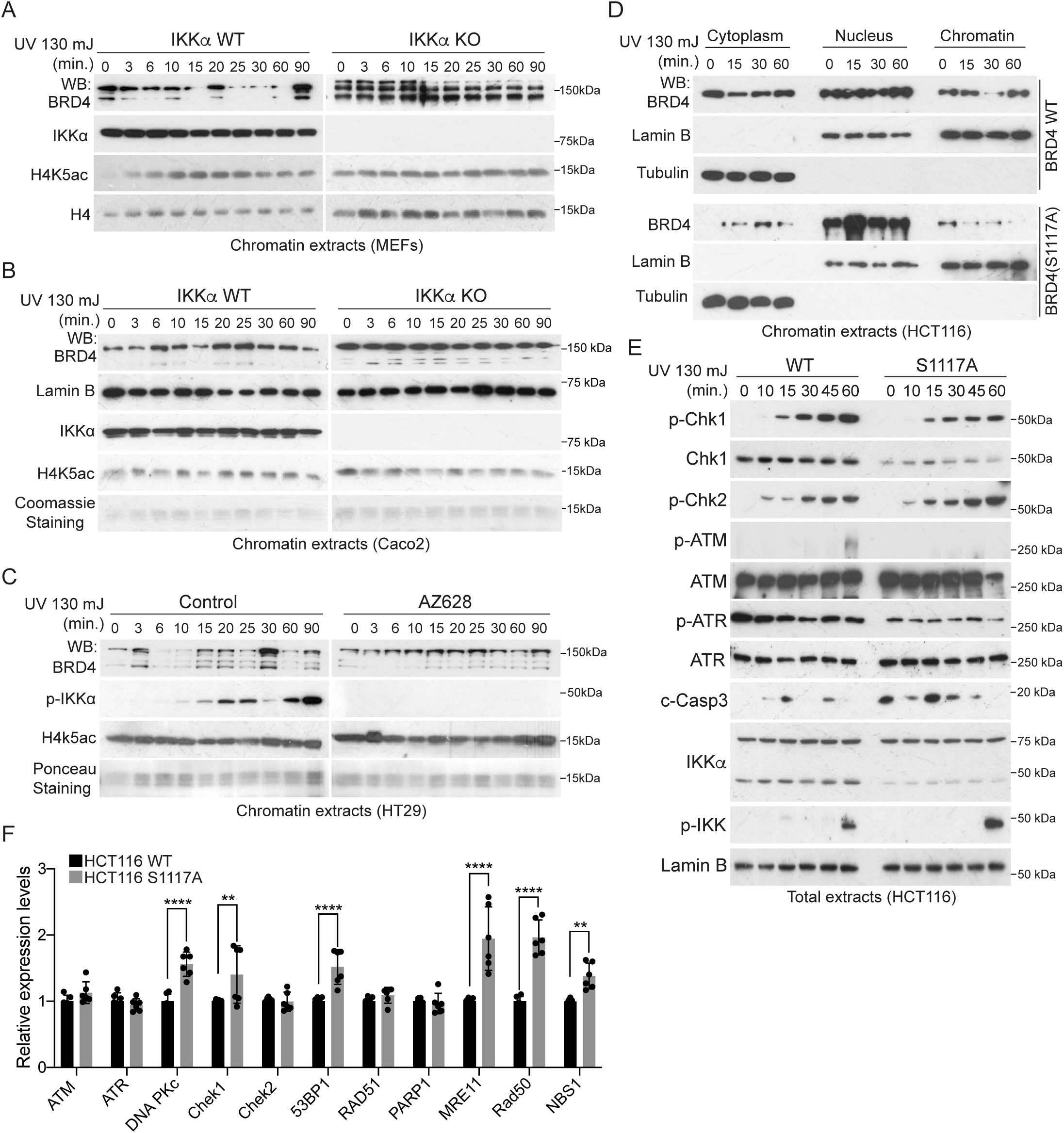
IKKα-induced phosphorylation regulates BRD4 chromatin-binding dynamics. **(A, B)** Western Blot analysis (WB) of chromatin extracts from MEFs (A) and Caco2 (B) IKKα WT and IKKα KO exposed to UV light (130 mJ) and collected at the indicated time points (from one out of three independent experiments). **(C)** WB analysis of chromatin extracts from HT29 cell line treated with BRAF inhibitor (AZ628, 10 nM) for 16h before UV light exposure (130 mJ) and collected as indicated time points (from one out of three independent experiments). **(D)** HCT116 BRD4 WT and HCT116 BRD4_S1117A_ knock-in cells exposed to UV light (130 mJ) at the indicated time points followed by subcellular fractionation (cytoplasm, nucleus and chromatin) and WB analysis (from one out of three independent experiments). **(E)** Western blot analysis of HCT116 BRD4 WT and HCT116 BRD4_S1117A_ knock-in cells treated as indicated (from one out of three independent experiments). **(F)** Expression analysis by qPCR of the indicated DDR-related genes from WT and BRD4_S1117A_ knock-in HCT116 cells. Bars represent mean values and standard error of the mean (s.e.m.) from 6 independent replicates; p-values were derived from two-sided unpaired T-test. ****p-value < 0.0001, **p-value <0.0025.

### IKKα dictates BRD4 chromatin binding to specific STAT3 target genes

To investigate whether aberrant chromatin-binding dynamics in IKKα deficient cells led to different genomic BRD4 occupancy after damage, we took advantage of the murine organoid model derived from APC^Min/+^ IKKα WT and KO intestinal tumors^33^. We first confirmed that S1117 of BRD4 was phosphorylated in APC^Min/+^ intestinal organoids (Figure S3A) and IKKα deficiency precluded chromatin binding dynamics of BRD4 upon IR (Figure S3B). We performed ChIP assay with BRD4 antibody from IKKα WT and KO APC^Min/+^ organoids left untreated or treated with IR and recovered after 60 minutes. We observed a comparable distribution of BRD4 at the different genomic regions (intergenic, promoter or gene body) between IKKα WT and KO cells both in untreated conditions and after IR (Figure S3C). The read coverage extracted from ChIP-seq experiments across 1kb non-overlapping genomic windows showed a significant correlation between WT and IKKα KO cells in non-irradiated conditions (Figure 4A), indicating that basal BRD4 binding landscape was IKKα independent. However, IR treatment resulted in the detection in IKKα WT cells of a subset of BRD4-bound regions corresponding to 774 genes (Figure 4B, red dots) that were marginally detected in IKKα deficient cells (Figure 4C, purple dots). These results indicated that a number of IR-induced BRD4 binding sites were IKKα dependent, which was further supported by correlation analysis of IR-treated IKKα WT and KO cells (purple dots in Figure 4D). Next, we performed a transcription factor (TF) target gene enrichment analysis of regions bound by BRD4 in the different experimental conditions. In non-irradiated conditions BRD4-bound regions showed a significant representation of target genes for multiple transcription factors including MYC, UBTF, GATA2 or RELA (Figure 4E). Interestingly, BRD4-bound genes exclusively detected in IR-treated IKKα WT cells were specifically enriched in UBTF, TRIM28 and STAT3 target genes (Figure 4E). We observed a significant correlation between IKKα- and IR-dependent BRD4-bound genes (Figure 4F) with STAT3 target genes^34,35^ (Figure 4G and Supplementary Table S1). Among the STAT3 target genes recruiting BRD4 in an IKKα-dependent manner, we identified *Bcl3*^36^ (Supplementary Table S1 and Figure S3D), which is an essential inhibitor of apoptosis in several systems^37,38^ and facilitates immune scape in cancer through PDL1 upregulation^39^. By qPCR, we found that basal and IR-induced *Bcl3* expression was decreased in IKKα-deficient APC^Min/+^ organoids (Figure 4H) and MEF cells (Figure 4I). *Bcl3* levels were also reduced in cells treated with the BRD4 inhibitor JQ1 (Figure 4J) and the JAK/STAT inhibitor ruxolitinib (Figure 4K), indicating that *Bcl3* expression requires simultaneous BRD4 and STAT3 activity. We confirmed altered IR-induced expression of several double BRD4 and STAT3 targets by qPCR analysis of APC^Min/+^ IKKα KO organoids (Figure S3E).

**Figure 4.**
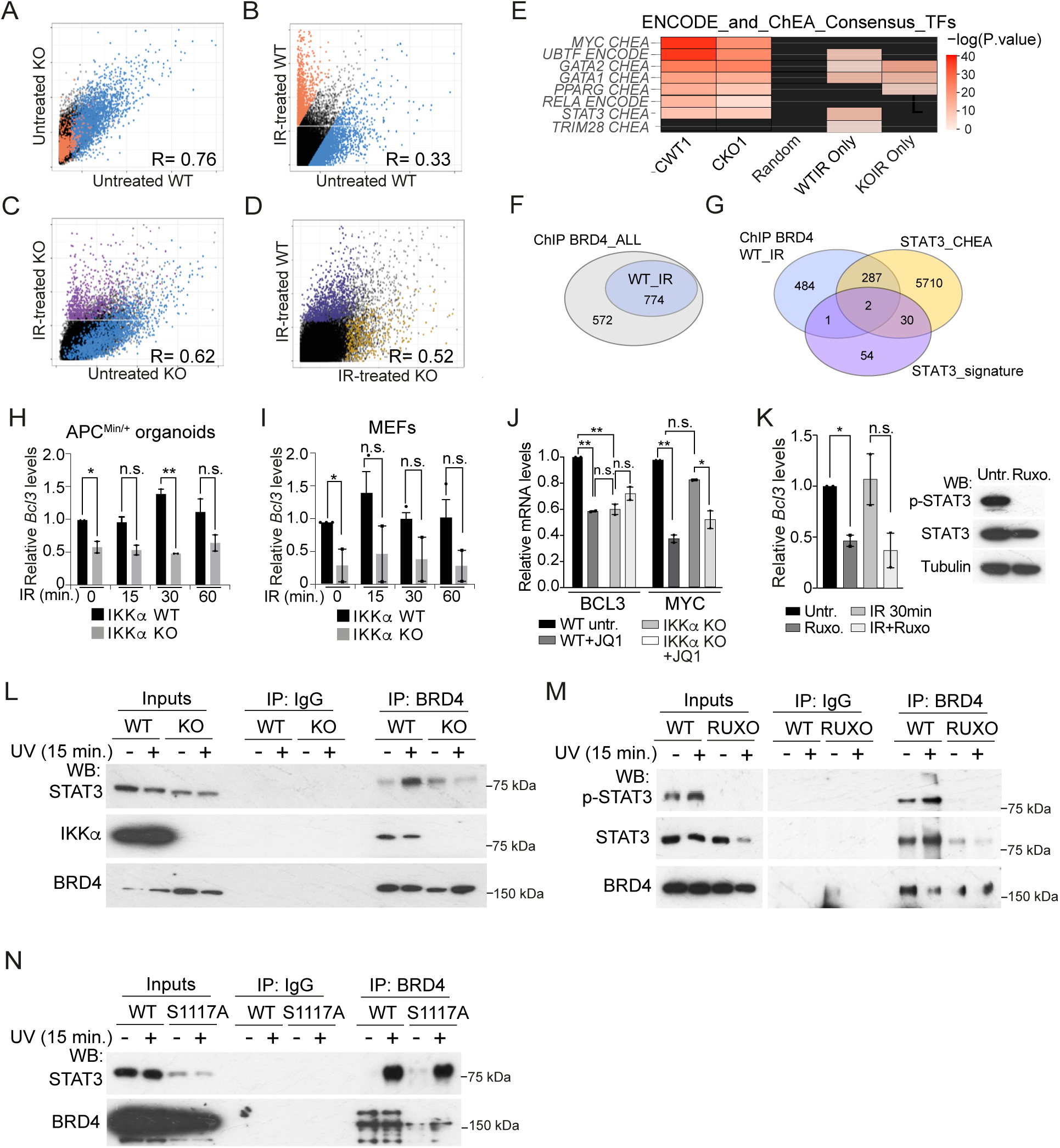
IKKα dictates BRD4 chromatin binding to specific STAT3 target genes. (**A-D**) Correlation of ChIP-seq depth across the genome of untreated or IR-treated WT and IKKα KO APC^Min/+^ organoid samples. Untreated IKKα WT and KO (A), IR-treated and untreated IKKα WT (B), IR-treated and untreated IKKα KO (C) and IR-treated IKKα WT and KO (D). In all the plots dots correspond to 1 kb non-overlapping genome windows. Dots in blue represent genomic regions that show a higher coverage in untreated WT compared with IR-treated WT, according to a linear model. In red are regions that show a higher coverage in IR-treated WT compared with untreated WT. In purple, regions with a higher coverage in IR-treated KO compared to untreated KO; and in yellow, regions with a higher coverage in IR-treated IKKα KO compared to IR-treated WT. **(E)** Enrichment analysis of different TFs targets from ENCODE and CHEA databases within each group of BRD4-bound regions (CWT, regions specifically detected in the WT untreated; CKO, regions detected in untreated IKKα KO; “random” refers to randomly selected regions of the genome; WTIR, regions exclusively detected in IR-treated IKKα WT; and KOIR regions exclusively found in IR-treated IKKα KO cells). **(F, G)** Venn diagrams representing the number of BRD4-bound genes upon IR treatment that are IKKα dependent (F) and those who have been previously identified as STAT3 targets (G). **(H-K)** qPCR gene expression analysis of the double BRD4 and STAT3 target gene *Bcl3* in APC^Min/+^ organoids (H, J and K) or MEFs (I) treated as indicated (IR; 10 Gy). Reduced levels of MYC (J) and p-STAT3 (K) indicated the efficacy of the treatments. Bars in H-K represent mean values and standard error of the mean (s.e.m.) from two independent replicates; p-values were derived from two-sided unpaired T-test in H, I and K and from one-way ANOVA in J. **p-value <0.0025, *p-value < 0.05; n.s.: no significant. In L-N, inputs correspond to 1/25 of the precipitated lysate. **(L, M)** IP with anti-BRD4 antibody from untreated or UV-treated IKKα WT and KO APC^Min/+^ organoids (L) or IKKα WT MEFs untreated or treated with 10 µM ruxolitinib (Rux) for 16 h and then exposed to UV light (130 mJ) for 15 min (M). (from one out of three independent experiments). **(N)** IP with anti-BRD4 antibody from parental and HCT116 S1117A knock-in HCT116 cells collected 15 min. after UV exposure (from one out of three independent experiments).

These results suggested that IKKα was coordinating BRD4- and STAT3-dependent transcription upon damage. Supporting this concept, we detected basal BRD4 and STAT3 interaction by CoIP, which was increased after UV treatment and abrogated in IKKα KO cells (Figure 4L). Basal and UV-induced association between BRD4 with STAT3 was abolished by the JAK/STAT inhibitor ruxolitinib (Figure 4M) but unaffected by the BRD4 S1117 mutation (Figure 4N). Together these results indicate that BRD4 preferentially binds to active STAT3, which is IKKα dependent. However, BRD4 and STAT3 interaction is independent of BRD4 phosphorylation by IKKα.

### IKKα deficiency precludes STAT3 activation in the presence of a functional JAK/STAT signalosome

We studied the possibility that IKKα kinase was regulating STAT3 activation upon damage. In agreement with this possibility, we detected basal and damage-induced phosphorylation of STAT3 at tyrosine 705 (target of the Janus kinase, JAK, and marker of canonical STAT3 activation) in IKKα WT cells, which was primarily abolished in cells lacking IKKα (Figures 5A-C, S4A-C). However, STAT3 activation was only marginally decreased after inhibition of nuclear IKKα(p45) by AZ628 ^5,40^ (Figure S4D), suggesting that was canonical IKK activity that regulates JAK/STAT. Since IKKs does not possess tyrosine kinase activity, we reasoned that IKKα may regulate the expression or activity of one or more upstream elements of the JAK/STAT signaling pathway. By qPCR analysis, we found that mRNA levels of *JAK1*, *JAK2*, *IL6R, LIFR* and the *GP130* co-receptor were comparable or higher in IKKα KO MEFs than in WT cells (Figure 5D). Supporting the functional integrity of the JAK/STAT signalosome, levels of p-STAT3(Y705) were robustly induced by treating IKKα KO MEFs with the canonical JAK/STAT activators IL6 and Leukemia Inhibitory Factor (LIF) (Figure 5E). Conditioned media (CM) from UV-treated WT MEFs also induced STAT3 activation in IKKα KO MEFs at levels comparable to IL6 or LIF (Figure 5E). Moreover, treatment of IKKα KO MEFs with LIF was sufficient to induce *BCL3* transcription, which was not observed upon IL6 treatment (Figure 5F). Together these results indicate that IKKα contributes to STAT3 activity through the regulation of one or more secreted JAK/STAT activating factors.

**Figure 5.**
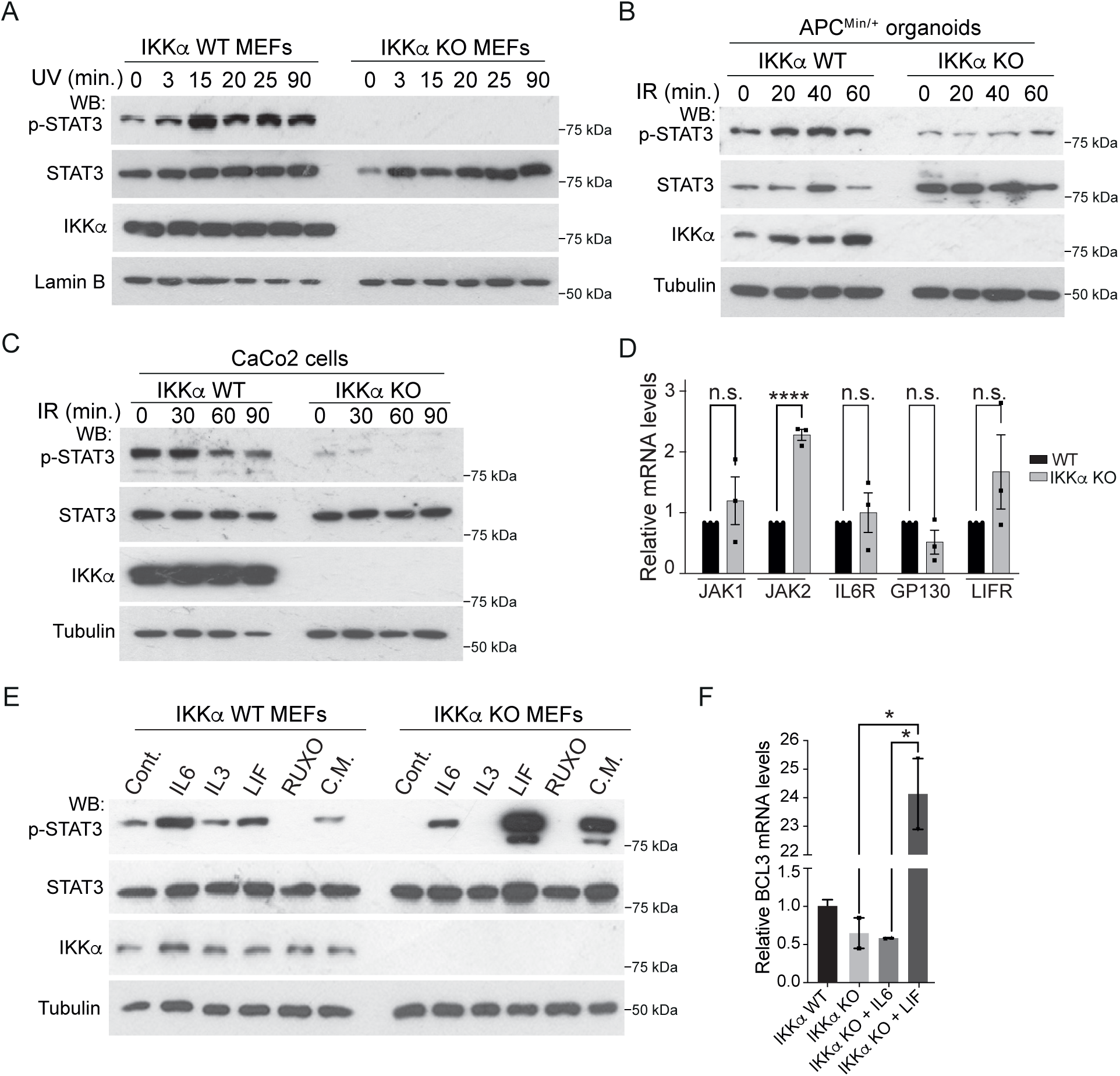
IKKα regulates STAT3 signaling by its paracrine activation through LIF. **(A-C)**WB analysis of extracts from WT and IKKα KO MEFs (A), APC^Min/+^ organoids (B) and Caco2 cells (C) exposed to ionizing radiation (IR) (10 Gy) and collected at the indicated time points (from one out of three independent experiments). **(D)** qPCR analysis of the indicated genes in MEFs IKKα WT and IKKα KO. **(E)** WB analysis of extracts from MEFs IKKα WT and KO pretreated with IL-6 (50 ng/ml), IL-3 (100 µg/ml), LIF (10 ng/ml) (Leukemia inhibitory factor) or ruxolitinib (Rux, 2.5 µM) for 16 h, or conditioned media (C.M.) from UV-treated (30 min.) WT MEFs (from one out of two independent experiments). **(F)** qPCR analysis of WT and IKKα KO cells untreated or treated with IL-6 (50 ng/ml) or LIF (10 ng/ml) for 16 h. Bars in D and F represent the mean value and standard error of three and two independent replicates performed, respectively. p-values were derived from two-sided unpaired T-test in D and from one-way ANOVA in F. ****p-value < 0.0001, *p-value <0.05; n.s.: no significant.

### JAK/STAT and IKKα(p45) inhibition potentiates anticancer therapy and allow the effective eradication of CRC xenografts in vivo

The JAK/STAT pathway is a positive regulator of cell proliferation and inhibitor of apoptosis in multiple cellular systems (reviewed in^41^). We investigated the functional impact of IKKα depletion, which precludes STAT3 activation, in the apoptotic induction of CRC patient-derived organoid (PDO) cells upon CT treatment. By WB analysis, we found that IKKα-deficient PDO5 (Figure 6A) and PDO8 (Figure S5A) cells showed increased apoptosis after 5-FU+Iri treatment compared with parental cells as determined by cleaved-PARP1 detection. Supporting the involvement of JAK/STAT and BRD4 in the therapy resistance imposed by IKKα, treatment of PDO5 with ruxolitinib or JQ1 significantly increased 5-FU+Iri sensitivity in dose-response assays (p<0.0001, for both conditions, Two-way ANOVA test) (Figure 6B). Antitumor effects of CT were increased by AZ628, which has little impact on STAT3 activity (Figure S4D) but precludes IKKα-dependent activation of ATM^5^ and BRD4 (see Figure 3C), and further enhanced by addition of the STAT3 inhibitor ruxolitinib (Figures 6C and S5B) (5-FU+Iri, IC50=2.37 mg/ml; 5FU+Iri+AZ628, IC50=0.025 mg/ml and IC50=1.982e-006 mg/ml for the quadruple combination in PDO5. p<0.0001 for all comparisons, Two-way ANOVA test). In contrast, we did not detect any additional benefit of combining 5-FU+Iri and AZ628 with the BRD4 inhibitor JQ1 (Figure S5C) suggesting that AZ628 was already inhibiting IKKα-dependent BRD4 activity in PDO cells, similar to that found in HT29 cells (see Figure 3C). Then, we tested the therapeutic efficacy of combining STAT3 and IKKα(p45) inhibition with CT in vivo. We implanted equivalent volumes of a human *KRAS* mutated CRC tumor with partial resistance to 5-FU+Iri (CRC#3) in the cecum of athymic Nude Mice. Tumor growth was monitored by palpation and, at the time of tumor detection, mice were randomly ascribed to the treatment groups: Vehicle, 5-FU+Iri, 5-FU+Iri plus the BRAF inhibitor vemurafenib (Vem.), 5-FU+Iri plus the JAK/STAT inhibitor ruxolitinib (Rux) or the quadruple combination 5-FU+Iri+Vem+Rux. As previously demonstrated in PDO-derived tumors^5^, combination of 5-FU+Iri plus vemurafenib significantly reduced tumor growth compared with 5-FU+Iri treatment that was slightly increased by the addition of the JAK/STAT inhibitor ruxolitinib (Figures 6D-F). However, microscopic examination of the tumors demonstrated the presence of extensive areas of necrosis and fibrosis in the 5-FU+Iri+Vem and 5-FU+Iri+Rux groups, which practically occupied the whole tumor mass in the quadruple treatment group (Figure 6G, see pink areas surrounding the alive tumor territory, T), with residual neoplastic cells displaying a severe pleomorphism associated to the presence of pyknotic nuclei (Figure 6H). We detected a substantial inhibition of cell proliferation in the residual tumor areas of all tested combination treatments, which was more significant in the quadruple combination group (Figures 6I and 6J). Importantly quadruple combination treatment did not impose any measurable toxicity in the animals as indicated by the overall appearance of mice and the absence of histological or functional alterations (i.e. proliferation) in the colonic tissue adjacent to the implanted tumors (Figure S5D). We obtained largely comparable results in a second *in vivo* experiment using an *APC, KRAS* and *PI3K* mutant CRC pulmonary metastasis (MPL-CRC7) (Figures S5E-H), further indicating the robustness of the combination treatment.

**Figure 6.**
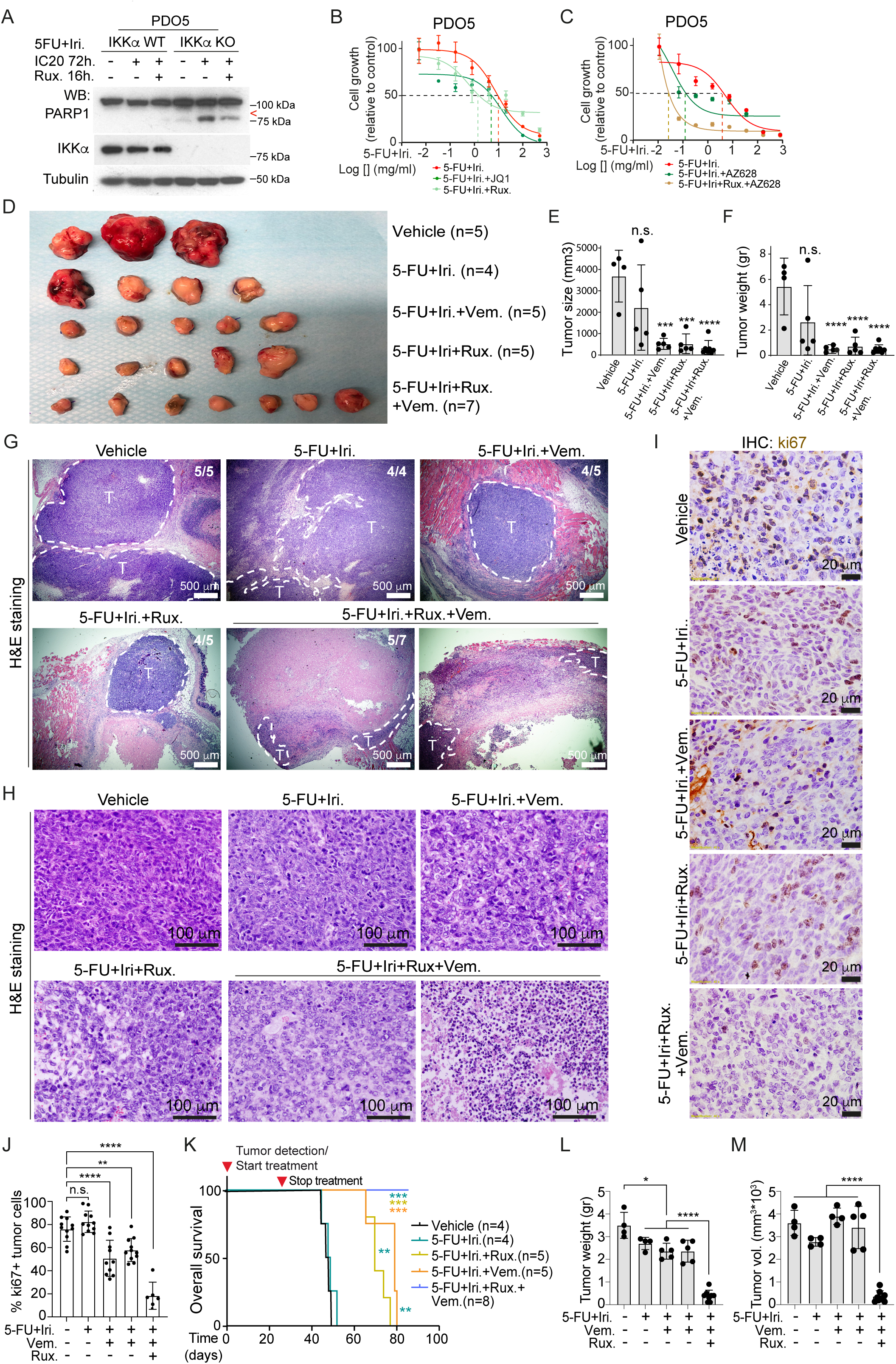
Addition of JAK/STAT and BRAF inhibitors to therapeutic treatment leads to effective eradication of chemotherapy-resistant human tumors. **(A)** WB analysis of IKKα WT and KO PDO5 cells treated with IC20 5-FU+Iri for 72 hours and ruxolitinib (Rux, 40 µM) for 16 hours (from one out of two independent experiments). **(B, C)** Dose-response curves of PDO5 cells treated with 5-FU+Iri alone or in combination with ruxolitinib [40 µM] (B) or 5-FU+Iri and AZ628 [25 µM] with or without ruxolitinib (C) for 72 hours (n = 3 replicates examined, from 1 out of 3 biologically independent experiments). 5-FU, 5-fluorouracil: Iri, irinotecan. **(D-F)** Photograph of tumors recovered from mice with xenografts in the indicated groups of treatment (D) and quantification of the size (E) and weight **(F)** of tumors. Of note that two of the tumors in the vehicle group were sacrificed during the weekend and processed by the personal in the animal facility, consequently they were not photographed and only one was measured. **(G, H)** Representative microscopic images of tumors recovered from mice in the indicated groups of treatment showing the areas containing alive tumor cells labelled as T and delimited by dashed lines (G). Detail of the residual tumor areas (H). Notice the presence of massive fibrotic and necrotic areas surrounding the tumor areas. Number of tumors displaying the exposed phenotype (X) from the total of animals in each group (N) is indicated in the upper-right corner of the images (X/N). **(I, J)** IHC analysis of the proliferation marker ki67 in the remaining tumor areas of the different groups of treatment (I) and quantification (J) of 10 areas per group (40X), when possible. **(K)** Kaplan-Meier curves for mice bearing a metastatic human tumor treated as indicated. Overall survival is indicated for the different treatments and statistical significance was determined using the Mantel-Cox log-rank test. In the legends Vem, vemurafenib; 5-FU, 5-fluorouracil; Rux, ruxolitinib and Iri, irinotecan. **(L, M)** Total weight (L) and volume (M) of tumors recovered at humanitarian endpoint for each mouse or at the end of the experiment in the case of quadruple combination treatment group. Bars in E, F, J and L-M represent mean values and standard deviation and statistical significance was determined by one-way ANOVA test and the Tukey’s Multiple Comparison Analysis. *p-value < 0.05, **p-value < 0.01, ***p-value < 0.001, ****p-value <0.0001; n.s.: no significant.

We then examined the long-term therapeutic potential of the quadruple combination of 5-FU+Iri+Vem+Rux in a third chemotherapy resistant CRC pulmonary metastasis carrying *APC*, *KRAS* and *TP53* mutations (MPL-CRC4). In agreement with the chemotherapy resistance observed in the patient, we did not observe any significant improvement in the survival of the 5-FU+Iri-treated group compared with the vehicle-treated control, which was clearly observed with the addition of vemurafenib or ruxolitinib (Figure 6K). As expected, we did not detect significant differences in tumor weight or volume at the time of sacrifice in these groups of treatment (Figures 6L and 6M). Importantly, none of the animals treated with the quadruple combination of 5-FU+Iri+Vem succumbed in the period of the study as shown in the Kaplan-Meier curves (Figure 6K). Importantly, and further indicating the high efficacy of combining 5-FU, irinotecan and vemurafenib, tumors present in this group of mice at the experimental endpoint were largely reduced compared with all other groups, which had been sacrificed earlier (Figure 6L and 6M). These results indicate that combination of BRAF and JAK/STAT inhibitors with DNA-damaging agents could provide a long-term therapeutic benefit to patients carrying tumors and/or metastases with different mutational burden that have acquired resistance to first-line chemotherapy treatment.

### LIF mediates JAK/STAT activation by IKKα thus imposing therapeutic resistance in CRC cells and poor patient outcome

We have previously identified the Leukemia Inhibitory Factor (LIF) as a crucial mediator of JAK/STAT activation^27^ and immunotherapy refraction^28^ in cancer. We investigated the possibility that LIF was inducing JAK/STAT signaling downstream of IKKα. By qPCR analysis, we detected variable levels of LIF, LIFR and GP130 mRNA in all tested cellular models (Figure S6A). Importantly, LIF mRNA levels were consistently reduced in APC^Min/+,^ MEFs, Caco2 cells and PDO5 following IKKα depletion (Figure 7A). By ChIP assay, we detected p65/NF-κB binding to the promoter region of the murine *LIF* gene specifically in the IKKα WT cells (Figure 7B) strongly suggesting that LIF transcription was induced by canonical NF-κB, downstream of IKKα.

**Figure 7.**
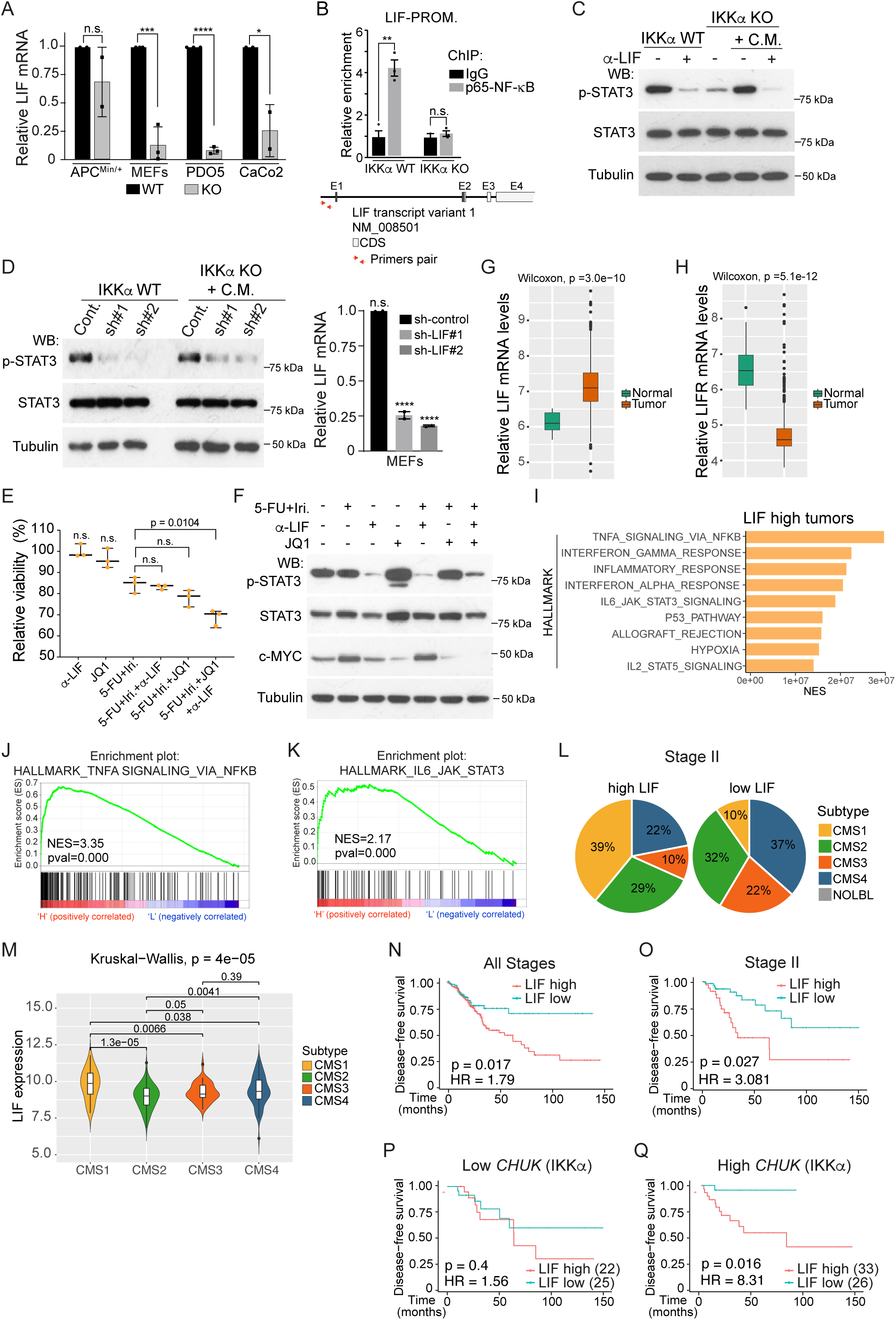
LIF mediates JAK/STAT activation by IKKα thus imposing therapeutic resistance in CRC cells and poor patient outcome. **(A)** qPCR analysis of LIF levels in the indicated IKKα WT and KO cells (n = 2 (APCmin) and 3 (MEFs, PDO5, Caco2) biologically independent experiments). **(B)** ChIP-qPCR analysis of p65/NF-κB binding to the promoter region of the murine LIF gene in MEFs IKKα WT and KO (n = 3 biologically independent experiments). **(C)** WB analysis of untreated and anti-LIF-treated (100 µg/ml for 2 hours) IKKα WT MEFs and IKKα KO MEFs incubated for 2 hours with control or anti-LIF-treated IKKα WT conditioned media (C.M.) (from one out of two independent experiments). **(D)** WB analysis (left panel) of control and LIF shRNA-transduced IKKα WT MEFs and IKKα KO incubated with control medium or conditioned media from the indicated cells treated with IR for 30 min (from one out of two independent experiments). qPCR analysis of LIF levels (right panel) of control and LIF shRNA-transduced IKKα WT and IKKα KO MEFs (n = 2 biologically independent experiments). **(E, F)** Survival assay in HCT116 CRC cells treated as indicated (E) and WB analysis of the same cells with the indicated antibodies (F) to evaluate the effects of the α-LIF antibody and JQ1 on p-STAT3 and MYC levels, respectively (n = 3 biologically independent experiments). **(G, H)** Box plots of LIF **(G)** and LIFR **(H)** levels in tumors samples (n=566) compared with normal samples (n=19) from the Marisa dataset. Boxes represent the central 50% of the data (from the lower 25th percentile to the upper 75th percentile), lines inside boxes represent the median (50^th^percentile), and whiskers are extended to the most extreme data point. Statistical p-value from Wilcoxon two-sided test is shown. **(I)** Barplot depicting the normalized enrichment score of statistically significant enriched pathways obtained by GSEA analysis with the Hallmark gene set for patients with high-LIF expression (NOM p-val<0.05). **(J, K)** GSEA of the TNFα via NF-κB (B) and JAK/STAT (C) pathway in high-LIF (n=159) versus low-LIF (n=170) tumors in TCGA patients. p-value is nominal p-value given by the GSEA program in J and K. **(L)** Pie charts showing the molecular subtype distribution according to Guinney et al^44^ in patients carrying LIF high (n=55) and LIF low (n=51) tumors from the stage II TCGA cohort. From 55 high LIF tumors, 41 with CMS information: CMS1 n=16, CMS2 n=12, CMS3 n=4 and CMS4 n=9. From 51 low LIF tumors, 41 with CMS information: CMS1 n=4, CMS2 n=13, CMS3 n=9 and CMS4 n=15. **(M)** Graphical representation of LIF levels in tumors corresponding to the different CMS subtypes. Statistical p-value from Kruskal-Wallis test is shown. **(N, O)** Kaplan-Meier representation of disease-free survival (DFS) over time for all patients in the TCGA CRC cohort (E) (LIF high n=222 and LIF low n=107) and selected stage II patients (F) (LIG high n=37 and LIF low n=37) according to LIF mRNA levels (optimized cutoff of 8.94 and 9.89, respectively). **(P, Q)** Kaplan-Meier representation of disease-free survival (DFS) over time for Stage II patients in the indicated groups according to *CHUK* and *LIF* levels. In A, B, D and E, bars represent mean values and standard error of the mean (s.e.m.). p-values were derived from two-sided unpaired T-test in A and B and from one-way ANOVA in D and E. ****p-value < 0.0001, ***p-value < 0.001, **p-value <0.01 *p-value < 0.05; n.s.: no significant. In N-Q, the optimized cut-off for LIF-high and low was determined using the ‘maxstat’ R package. In N-Q, we used Cox proportional hazards models for statistical Kaplan–Meier analysis and log-rank two-sided p-value. HR, hazard ratio.

Then, we tested whether STAT3 activity was affected upon abrogation of LIF activity with the anti-LIF blocking antibody (Figure 7C) or with two specific shRNAs (Figure 7D) in IKKα proficient cells. LIF inhibition by anti-LIF treatment or by shRNA led to a reduction in p-STAT3 (Y705) levels in IKKα WT MEFs and precluded the capacity of conditioned media to induce STAT3 activation in IKKα KO cells (Figures 7C and 7D).

We then tested whether inhibiting LIF and BRD4 improved the efficacy of chemotherapy in CRC cells. We treated HCT116 cells with sublethal doses of 5-FU+Iri alone or in combination with the BRD4 inhibitor JQ1, anti-LIF blocking antibody or both. Adding JQ1 or anti-LIF antibody to the chemotherapy treatment did not result in a significant improvement of chemotherapy efficacy, which was reproducibly and significantly observed in the triple combination (Figure 7E). By WB assay, we found that anti-LIF and JQ1 treatments led a significant reduction in the levels of p-STAT3 and the BRD4 target c-MYC^42^, respectively (Figure 7F).

Finally, we investigated whether the IKKα-LIF-JAK/STAT axis had any impact in CRC patients. We found that LIF was expressed in all tumor samples from Marisa^43^ and TCGA (The TCGA Portal) datasets with variable expression levels (Figures 7G and S6B). LIF levels were significantly higher in tumors compared with normal adjacent mucosa (Figure 7G), whereas LIFR show an inverse correlation (Figure 7H). Gene Set Enrichment Analysis (GSEA) of expression data from the TCGA cohort uncovered TNFα via NF-κB as the main activated pathway in high *LIF* CRC tumors when compared with CRC tumors with low *LIF* mRNA (Figure 7I and 7J) followed by Interferon response and the JAK/STAT pathway (Figure 7I and 7K).

*LIF* high tumors were primarily ascribed to the CMS1 subtype, in stage II CRC patients, according to the Guinney classification^44^ (Figure 7L), which comprises the majority of tumors carrying microsatellite instability (MSI). The CMS1 is characterized by the presence of a diffuse immune infiltrate but a paradoxical activation of immune evasion pathways, which is a characteristic of MSI tumors^45^. Accordingly, LIF mRNA levels were significantly higher in the group of MSI-related tumors (CMS1) compared with the rest of tumor subtypes (Figure 7M). Analysis of the differentially expressed genes (DEG) in the *LIF* high tumors identified a significant enrichment in functions related to immune response and T cell activity (Figure S6C) positioning this type of tumors as candidates for immunotherapy treatment likely in combination with LIF inhibitors. Further bioinformatic analysis demonstrated that *LIF* levels were sufficient to stratify patients displaying the poorest disease-free-survival both at all stages (p=0.017; HR=1.79) (Figure 7N) and in the subset of stage II patients (p=0.0027; HR=3.08) (Figure 7O), which is particularly relevant in the clinical context. Then, we studied the possible correlation between *CHUK* (the gene codifying for IKKα) and *LIF* levels in CRC. We observed that *LIF* high and low tumors were similarly distributed in the *CHUK* high and low groups (Figure S6D) with no significant correlation between *CHUK* and *LIF* mRNA levels (Figure S6E and S6F), suggesting that *LIF* expression can be regulated independently of IKKα in human tumors. Importantly, the prognosis value of *LIF* was restricted to tumors carrying high *CHUK* expression (Figures 7P and 7Q), being patients with *CHUK* and *LIF* high tumors the ones with the worst disease-free survival (HR=8.31, p=0.016) (Figure 7Q). These results suggested that pro-tumorigenic activity of LIF is sustained by additional IKKα activities such as BRD4 or ATM regulation and postulate LIF and IKKα/NF-κB as candidate prescription biomarkers for immunotherapy.

Together our data support a model in which IKKα imposes therapeutic refraction in CRC cells at different levels. On the one hand, IKKα kinase activity facilitates DNA repair downstream of ATM^5^ and BRD4^9^ (see model in Figure 8). On the other hand, IKKα through NF-κB promotes *LIF* transcription and JAK/STAT pathway activation to prevent early apoptosis upon DNA damage and to facilitate STAT3/BRD4 association. Because high *LIF* levels are largely detected in the subset of CMS1 (MSI) tumors and its prognosis value is linked to the presence of high *CHUK* mRNA, we propose *CHUK*/IKKα and *LIF* as potential biomarkers of immune evasion in MSI CRC, which should be further investigated in the context of immunotherapy.

**Figure 8.**
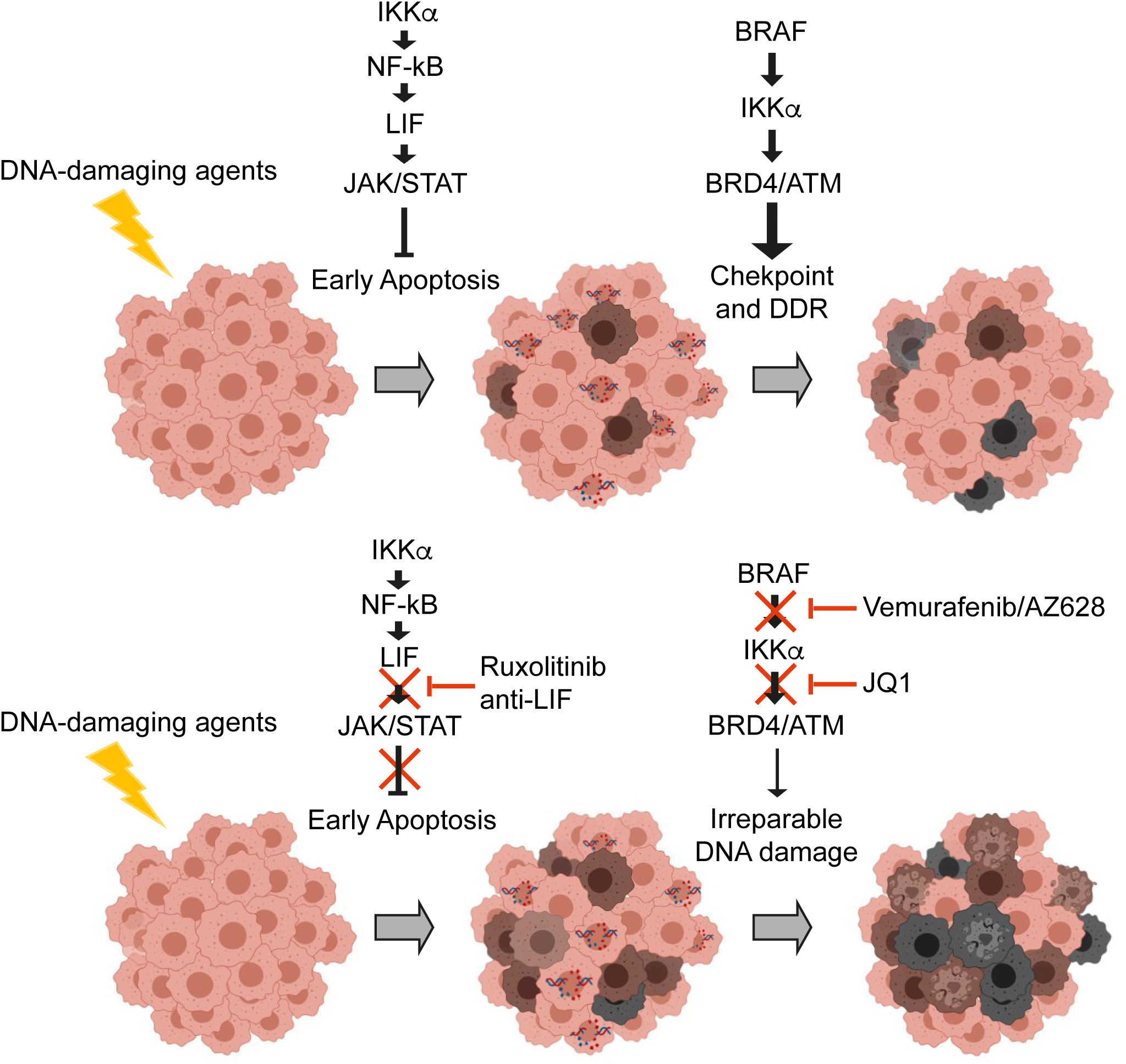
Model for IKKa-mediated therapy resistance promotion. In response to DNA damaging treatment, few cancer cells enter apoptosis (brown cells), which is partially prevented by STAT3 activation downstream of IKKα, whereas other cells accumulate variable amounts of DNA damage that is counteracted by activation of IKKα- and ATM-dependent DDR pathway. Inhibition of STAT3 by ruxolitinib or anti-LIF antibody and the DDR pathway by BRAF inhibitor or JQ1 (or abrogation of both pathways by IKKα depletion) results in massive cell death after CT treatment (dead cells are marked in dark grey).

## DISCUSSION

We had previously demonstrated that IKKα is an upstream regulator of the DDR pathway and cells carrying active IKK display higher resistance to CT agents linked to a more efficient DNA repair^5^. We now show that IKKα also provides CT refraction by inhibiting apoptosis, previous to DNA repair, downstream of BRD4 and STAT3 activation. Notably, these IKKα functions are mainly NF-κB independent but dependent on the kinase activity of IKKα on specific phosphorylation substrates. We have now shown that IKKα is an upstream kinase of the BET protein BRD4 at S1117, which regulates BRD4 capacity for dissociation and reassociation to chromatin. This result is specifically relevant since S1117 is located at C-terminal domain (CTD) of BRD4, which functions as a transcriptional coactivator domain by binding the transcription elongation factor (p-TEFb) complex that regulates RNA polymerase (Pol II) activity and controls productive transcription elongation ^46^. Thus, the different cycling dynamics of BRD4 observed across cell lines and damaging agents may reflect differences in the levels and dynamics of transcriptional coregulators such as transcription factors, activated STAT3 or polymerase II, or in the patterns of histone H3 and H4 acetylation. Moreover, comparison of the BRD4 sequences across species demonstrated a complete conservation of S1117 residue thus reinforcing the idea that S1117 phosphorylation may impact on BRD4 activity. This possibility opens a new avenue of research focused on the cooperative contribution of BRD4 and IKKα to the DNA damage response, with putative implications in cancer therapy and in the design of novel treatment protocols. To our view, the broad spectrum of IKKα activities, other than NF-κB activation, points out this kinase as a preferential target for therapy in combination with DNA damaging agents as we previously proposed^5^. Using the intestinal adenoma model of APC^Min/+^ murine organoids, we uncovered a novel association between BRD4 and STAT3 at chromatin. Although this is not the best model for studying human CRC (more similar to the human familial adenomatous polyposis adenoma model), the functional link between BRD4 and STAT3 downstream of IKKα has been further validated in several cellular systems in this work. Moreover, because proinflammatory cytokines are canonical upstream activators of IKK, we anticipate that the mechanism of therapeutic refraction imposed by the IKKα/JAK/STAT/BRD4 and the IKKα/ATM axes could be specifically relevant in tumors with high contribution of the immune system (such as tumors with high microsatellite instability, MSI), and IKKα-based therapy could be even enhanced following immunotherapy treatment (i.e. inhibitors of PD1/PDL1). In this sense, we demonstrated that conditioned media from IKKα WT MEFs imposes a robust activation of STAT3 in IKKα deficient cells suggesting that factors produced by the stroma could be inducing resistance pathways in the adjacent tumor cells. Thus, we propose that targeting IKK or JAK/STAT3 will be particularly useful for potentiating the effect of immunotherapy protocols such as combinations of chemotherapy plus PD1 or PDL1 inhibitors that are currently being used for treating MSI high CRC tumors^47^.

Physical association between IKKα and STAT3 was previously found to impose IKKα stabilization and NF-kB activation by preventing its association to ubiquitin ligases^48^. Now, we identified BRD4 and STAT3 as direct downstream effectors of the IKKα kinase, which represents additional evidence of its prominent role in cancer, being BRD4 and STAT3 well-known tumor and therapy resistance drivers. Importantly, STAT3 activation by IKKα is imposed by paracrine activation of the upstream JAK kinases by LIF. Thus, it is likely that other important factors downstream of LIF such as STAT1 or elements of the MAPK or PI3K pathway^49^ could also contribute to the observed phenotype. Because LIF has recently been identified as a suppressor of anti-PD1 therapy^28,29^ and we found a clear association between LIF levels in CRC with MSI tumors and the immune-evasion phenotype, we propose that inhibitors of the IKKα kinase activity could have particular impact in tumors that are candidate for immunotherapy treatment. In the absence of safety and highly specific IKKα inhibitors that have already been approved for human therapy (reviewed in^50^), we propose that combination treatments involving CT plus BRAF inhibitors and/or inhibitors of the JAK/STAT and BRD4 pathways will represent a suitable strategy to revert chemo-refraction, thus leading to more efficient therapeutic protocols.

## AUTHOR CONTRIBUTIONS

L.E. and A.B. conceptualized the study, designed the experiments and wrote the manuscript. E.B. and E.S. designed proteomic analysis and evaluated results. I.P., L.S., A.V., D.A-V., T.L-J., J.A-M, J.B., Y.G., A.M., M.M-I., V.G-H., G.G., R.S., C.S. and M.G. performed biochemical assays, and *in vitro* and *in vivo* drug testing experiments. E.B-T, R.I. and J.S. provided reagents, helped in writing the manuscript and with interpretation of results. M.I. performed the clinicopathological characterization of human tumors. I.P. and L.E. prepared the figures.

## ACKNOWLEDGEMENTS

We want to thank Espinosa’s and Bigas’ lab members for constructive discussions and suggestions. This work was funded by grants from Instituto de Salud Carlos III FEDER (PI22/00069, PI19/00318, PI19/01320 and PT20/00023), Generalitat de Catalunya 2017SGR135, Xarxa de Bancs de tumors sponsored by Pla Director d’Oncologia de Catalunya (XBTC) and Fundación Asociación Española contra el Cáncer (AECC). DA-V is a recipient of the FI20/00130 grant from Instituto de Salud Carlos III FEDER. TL-J is a recipient of the AECC postdoctoral grant POSTD21975. CRG/UPF Proteomics Unit is part of the “Plataforma de Recursos Biomoleculares y Bioinformáticos (ProteoRed)” supported by grant PT13/0001 of Instituto de Salud Carlos III from the Spanish Government and “Secretaria d’Universitats i Recerca del Departament d’Economia i Coneixement de la Generalitat de Catalunya” (2017SGR135).

## CONTACT FOR REAGENT AND RESOURCE SHARING

Further information and requests for resources and reagents should be directed to and will be fulfilled by the Lead Contact, Lluís Espinosa.

## EXPERIMENTAL MODEL AND SUBJECT DETAILS

### Patient-derived and mouse intestinal organoids

For patient-derived organoids (PDOs) generation, primary or xenografted human colorectal tumors were disaggregated in 1 mg/mL collagenase II (Sigma) and 20 µg/mL hyaluronidase (Sigma), filtered in 100 µm cell strainer, and seeded in Matrigel (BD Biosciences) as described ^51^. PDOs were expanded by serial passaging and kept frozen in liquid Nitrogen for being used in subsequent experiments. PDO5 is TP53 WT and carries the KRAS G12D mutation. PDO8 is TP53 Q192stop and KRASG13C. Samples from patients were kindly provided by MARBiobank, integrated in the Spanish Hospital Biobanks Network (RetBioH; www.redbiobancos.es). Informed consent was obtained from all participants and protocols were approved by institutional ethical committees.

APC^Min/+^-derived organoids from WT or IKKα KO mouse intestines were obtained as previously described ^33^.

### Cell lines

CRC cell lines HCT116 (*KRAS* mutated), Caco2 (*KRAS* and *BRAF* WT) and HT29 (*BRAF* mutated) were obtained from the American Type Culture Collection (ATCC, USA). WT and IKKα KO MEFs were kindly provided by Michael Karin (UCSD, La Jolla) and HeLa S3 cells were kindly provided by Dr. Shannon Lauberth, UCSD. All cells were grown in Dulbecco’s modified Eagle’s medium (Invitrogen) plus 10 % fetal bovine serum (Biological Industries) and were maintained in a 5 % CO_2_ incubator at 37 °C. Cells were routinely tested by PCR as being mycoplasma free.

IKK1 CRISPR knock out cells were generated by CRISPR-Cas9. Guides were designed using the online prediction tool created by Zhang lab at Massachusetts Institute of Technology (https://crispr.mit.edu). Two guides located between Exon 1 and Intron 1 on human *CHUK* gene (IKK1) with few predicted off targets were cloned into the LentiCRISPR V2 plasmid (Addgene plasmid#52961).

### Animal studies

To perform in vivo drug testing, equivalent pieces of individual tumors were implanted orthotopically in the wall of the cecum of athymic Nude Mice. When tumors were detectable by palpation (4-5 weeks), animals were randomly ascribed to the different groups of treatment. Vemurafenib (50mg/kg) and ruxolitinib (100mg/Kg) were administered orally every day, 5-FU (75mg/kg, divided in two doses) and irinotecan (20mg/Kg) every 4 days intravenously. After 21 days of treatment or at the humanitarian endpoint (in the case of the long-term survival experiment), mice were euthanized and tumors collected, photographed, measured and processed for immunohistochemistry examination. In all our procedures, animals were kept under pathogen-free conditions, and animal work was conducted according to the guidelines from the Animal Care Committee at the Generalitat de Catalunya. The Committee for Animal Experimentation at the Institute of Biomedical Research of Bellvitge (Barcelona) approved these studies.

### PDOs infection

sgRNA against *CHUK* gene was designed using Benchling. Lentiviral production was performed transfecting in HEK293T cells the lentiviral vectors and the plasmid of interest. One day after transfection, medium was changed, and viral particles were collected 24 hours later and then concentrated using Lenti-X Concentrator. PDOs were infected by resuspending single cells in concentrated virus diluted in complete medium, centrifuged for 1 h at 650 rcf, and incubated for 5 hours at 37°C. Cells were then washed in complete culture medium and seeded as described above.

### PDO viability assays

600 single PDO cells were plated in 96-well plates in Matrigel. After 6 days in culture, we treated growing PDOs with 5-FU, irinotecan, AZ628, ruxolitinib or combinations for 72 hours at the indicated concentrations. Cell viability was determined using the CellTiter-Glo 3D Cell Viability Assay (Promega) following manufacturer’s instructions in an Orion II multiplate luminometer (Berthold detection systems). Data were calculated as mean ± standard deviation, representing triplicates of one out of 2 independent experiments.

### Cell lysis and Western Blot (WB)

Cells were lysed 20 min at 4°C in 300 μl of PBS plus 0.5% Triton X-100, 1 mM EDTA, 100 mM NA-orthovanadate, 0.25 mM phenylmethylsulfonyl fluoride, and complete protease inhibitor cocktail (Roche). Lysates were analyzed by Western blotting using standard SDS–polyacrylamide gel electrophoresis (SDS-PAGE) techniques. In brief, protein samples were boiled in Laemmli buffer, run in polyacrylamide gels, and transferred onto polyvinylidene difluoride membranes. The membranes were incubated overnight at 4°C with the appropriate primary antibodies. After being washed, the membranes were incubated with specific secondary horseradish peroxidase–linked antibodies from Dako and visualized using the enhanced chemiluminescence reagent from Amersham. Primary antibodies used are listed in Supplementary Table S4.

### Cell fractionation

For cytoplasm/nuclear/chromatin separations, cells were lysed in 10 mM Hepes, 1.5 mM MgCl_2_, 10 mM KCl, and 0.05 % NP-40 (pH 7.9) for 10 min on ice and centrifuged at 3,000 rpm. Supernatants were recovered as the cytoplasmic fraction, and the pellets were lysed in 5 mM Hepes, 1.5 mM MgCl_2_, 0.2 mM EDTA, 0.5 mM dithiothreitol, and 26% glycerol and sonicated for 5 min three times to recover the soluble nuclear fractions. The remaining pellet included the chromatin fraction. Lysates were run in SDS-PAGE and transferred onto Immobilon-P transfer membranes (Millipore) for Western blot analysis.

### Immunoprecipitation assay (IP) and Pull-Down assay (PD)

PD assays were performed as previously described^52^. Briefly, GST fusion proteins were incubated with lysates for 45 min in a rotary shaker at 4°C. When indicated, nuclear extracts were boiled at 98°C for 5 min in the presence of 1% SDS to disassemble pre-existing protein complexes and then neutralized in 1% Triton X-100. Precipitates were resolved in SDS–PAGE and analyzed by IB. For peptide IP, histone H4 peptides [Synpeptide CO LTD] were synthesized as biotinylated N-terminal and C-terminal amides. Peptides were incubated overnight at 4°C with the indicated cell extracts and precipitated with streptavidin–sepharose beads for 45 min.

### Mass Spectrometry Analysis

Cell lysates obtained in the different experimental conditions were processed and digested with trypsin and endoproteinase LysC with a ratio enzyme:sample of 1:10 for both enzymes (w:w). Samples were then subjected to phospho-peptide enrichment using titanium dioxide (TiO_2_) beads, and phospho-enriched samples were analyzed by LC-MS/MS. To identify IKKα-dependent phosphopeptides, samples were injected with a 120-minute chromatographic gradient in an Orbitrap Velos Pro with a data-dependent acquisition method using CID fragmentation for the top 20 most intense precursor ions and multistage activation. In the UV-activation experiment, samples were acquired with a 90-minute gradient in an Orbitrap Fusion Lumos with a data-dependent acquisition method using top speed, HCD fragmentation and ion-trap detection. In both cases, the resulting data were analyzed with the Proteome Discoverer software v1.4, using the search algorithm Mascot (v2.5) against a Human protein database (Uniprot, v2015) with oxidation (Met), and phosphorylation (Ser, Thr, Tyr) as variable modifications. Carbamidomethylation (Cys) was set as fixed modification and a mass tolerance of 7 ppm (MS1) and 0.5 Da (MS2) were used. Only peptides with a false discovery rate below 5 % were considered for quantitative analysis. Peptides relative abundance was estimated with the area under the curve of extracted ion chromatograms. Protein network was generated using cytoscape software (www.cytoscape.org).

### ChIP sequencing and data Analysis

DNA samples were sequenced using Illumina HiSeq platform. Raw single-end 50-bp sequences were filtered by quality (Q >30) and length (length > 20 bp) with Trim Galore ^53^. Total filtered sequences, which ranged between 32 and 55 million per sample, were aligned against the reference genome (mm10 release) with Bowtie2 ^54^. MACS2 software ^55^ was run for each replicate considering unique alignments (q-value < 0.1). Peaks from biological replicates were merged using bedtools. Peaks that were detected in the input samples were filtered out, as well as the mouse black regions downloaded from the ENCODE portal (ENCFF547MET) ^56^ (https://www.encodeproject.org/). Peak annotation was performed with ChIPseeker ^57^ and Annotatr ^58^ packages; and functional enrichment analysis with enrichR ^59^, using the latest version of GO annotations. Peaks heatmaps and multiple coverage correlation across bigwig files were performed with Deeptools ^60^.

### Bioinformatics analysis

The transcriptome of normal and tumor samples from GSE39583 (Marisa et al) obtained by microarrays was downloaded from the Gene Expression Ommnibus (GEO) (Edgar R, Domrachev M, Lash AE.Gene Expression Omnibus: NCBI gene expression and hybridization array data repository Nucleic Acids Res. 2002 Jan 1;30(1):207-10) and analyzed with the affy R package. Transcriptomic and available clinical data from CRC TCGA dataset was downloaded from the open-access resource CANCERTOOL. We used the TCGA cohort (The TCGA Portal) to classified patients according to the mean expression of LIF. The association with relapse was assessed using Kaplan-Meier estimates and Cox proportional hazard models. A standard log-rank test was applied to assess significance between groups. This test was selected because it assumes the randomness of the possible censorship. All the survival analyses and graphs were performed with R using the survival (v.3.2-3) and survimer (v.0.4.8) packages and a p-value<0.05 was considered statistically significant.

Differentially expressed genes between patients with high versus down levels of LIF were explore using DESeq2 R package (v.1.24.0)^61^ from gene expression data downloaded from TCGAbiolinks R package^62^. Functional enrichment analysis was performed with enrichR, using Gene Ontology biological process annotation.

### Plasmids

Plasmid plentiCRISPRv2 was a gift from Feng Zhang (Addgene plasmid #52961; http://n2t.net/addgene:52961; RRID:Addgene_52961); plasmid p6344 pcDNA4-TO-Ha-Brd4FL, a gift from Peter Howley (Addgene plasmid #31351; http://n2t.net/addgene:31351; RRID:Addgene_31351); High fidelity Prime STAR HS DNA polymerase was from TAKARA. Oligonucleotides were purchased from sigma. The sequence of the different oligonucleotides used to generate the different constructs are listed in Supplementary Table S2.

### Plasmid construction

GST fusion proteins were generated using the vector pGEX5x.3. BRD4 and STAT3 coding sequences were cloned in frame downstream from GST, using BamHI and XhoI restriction sites. Sequences of interest were amplified by PCR using appropriate primers (Supplementary Table S2). Primers contain 5’ extensions to include restriction sites and to ensure the correct reading frame.

For STAT3, two fragments were fused to GST, amino acids 507 to 700 and 559 to 700. The STAT3 coding sequence was taken from Genebank, accession number NM_001369512. The starting material for cloning was cDNA prepared from HT29 cells RNA.

Several BRD4 fragments were produced as GST fusion proteins in *E. coli*. These include amino acids 2 to 320; 301 to 700; 685 to 1000, 1001 to 1362, 526 to 702, 577 to 641 and 1037 to 1201. To generate the appropriate plasmids, PCR was used on the template plasmid p6344 pcDNA4-TO-Ha-Brd4FL. The mutagenic primers used to generate the point mutation in codon 1117, that resulted in the substitution of serine with alanine, are depicted in Supplementary Table S2. Mutagenesis was performed by sequence overlap extension. For the full-length coding sequence, a fragment replacement was done taking advantage of a unique AgeI restriction site. All DNA constructs were verified by Sanger sequencing.

To modify the BRD4 gene *in situ*, two guide RNAs were designed using the Benchling platform (https://www.benchling.com/crispr/). One guide targets the double DNA cut 27 nucleotides 5’ of codon 1117 and the other 5 nucleotides 3’ of the stop codon. The information to produce these guide RNAs (primers gU and gD1117 in Supplementary Table S2) was introduced in plasmids pLentiCRISPR V2-BRD4.1 and pLentiCRISPR V2-BRD4.2 following published procedures ^63^. In addition, a repair plasmid, pBS-BRD4S1117A-ki, was constructed to achieve homologous recombination next to the targeted chromosomal locations. This plasmid carries two homology arms which extend 847 bp upstream and 906 bp downstream of the fragment to be replaced. The replacement DNA contains the desired codon 1117 mutation and several silent changes that rend the edited chromosome resistant non-recognizable by the guide RNAs. Moreover, this DNA fragment contains intronic and exonic sequences up to the BRD4 stop codon which is preceded by a loxP-P2A-mEmerald-loxP cassette. The mEmerald coding sequence was amplified from pmEmerald-Claudin7-C-12.

### CRISPR/Cas9-mediated BRD4 gene edition

HCT116 cells were co-transfected with the described targeting plasmids and the repair plasmid at a 1:1:40 molecular ratio. After 24 hours incubation, puromycin (1 μg/mL) was added for 72 hours to enrich for transfected cells and then removed. Cells were expanded for about two weeks and sorted and plated as single cells in 96-well plates to isolate clones.

### Kinase Assay

In vitro kinase assays were performed as previously described ^64^. In brief, 5 μg of GST or GST fusion proteins were incubated with 200 ng of recombinant human IKKα (ab102103) or 10 μg of cell lysates, as indicated, at 30° C for 30 minutes in the presence of ATP-γP^32^. Reactions were stopped by adding loading buffer, run in a polyacrylamide gel and developed in autoradiograph film. Primers used to generate GST fusion proteins are listed in Supplementary Table S2.

### RT-qPCR analysis

Total RNA from treated APC^Min/+^-derived organoids and MEFs cells was extracted with the RNeasy Micro Kit, and cDNA was produced with the RT-First Strand cDNA Synthesis Kit. RT-qPCR was performed in LightCycler 480 system using SYBR Green I Master Kit. Samples were normalized to the mean of the housekeeping genes *GAPDH* and *ACTB*. Primers used for RT-qPCR are listed in Supplementary Table S3.

### Hematoxylin and eosin staining

Previously de-paraffinized sections were incubated with hematoxylin 30 s, tap water 5 min, 80% ethanol 0.15% HCl 30 s, water 30 s, 30% ammonia water (NH3(aq)) 30 s, water 30 s, 96% ethanol 5 min, eosin 3 s, and absolute ethanol 1 min. Samples were dehydrated, mounted in DPX, and images were obtained with an Olympus BX61 microscope.

### Immunohistochemical staining

Paraffin blocks were obtained from tumor samples, previous fixation in 4% formaldehyde overnight at room temperature. Paraffin-embedded sections were de-paraffinized, rehydrated and endogenous peroxidase activity was quenched (20 min, 1.5% H_2_O_2_). Citrate based antigen retrieval was used. All primary antibodies were diluted in PBS containing 0.05% BSA, incubated overnight at 4 °C and developed with the Envision+ System HRP Labelled Polymer anti`-` Rabbit or anti`-`Mouse and 3,3′`-`diaminobenzidine (DAB). Samples were mounted in DPX and images were obtained with an Olympus BX61 microscope.

## QUANTIFICATION AND STATISTICAL ANALYSIS

Statistical parameters, including number of events quantified, standard deviation, and statistical significance are reported in the figures and in the figure legends. Statistical analysis has been performed using GraphPad Prism6 software (GraphPad) and p<0.05 is considered significant. Two-sided Student’s t test was used to compare differences between two groups and Two-Way ANOVA test was used to compare differences among multiple groups. Each experiment has been repeated at least twice.

## DATA AND SOFTWARE AVAILABILITY

MS data are available at PRIDE EBI-EMBL database with identifier PXD008932 and in ^5^.

ChIP-seq data for BRD4 is deposited at GEO (GSE196461) Token code: obepawuefrodlit

## SUPPLEMENTAL INFORMATION

Supplemental information includes 6 additional figures with their corresponding figure legends, and 4 supplementary tables.

---

**Supplementary Table S1:**
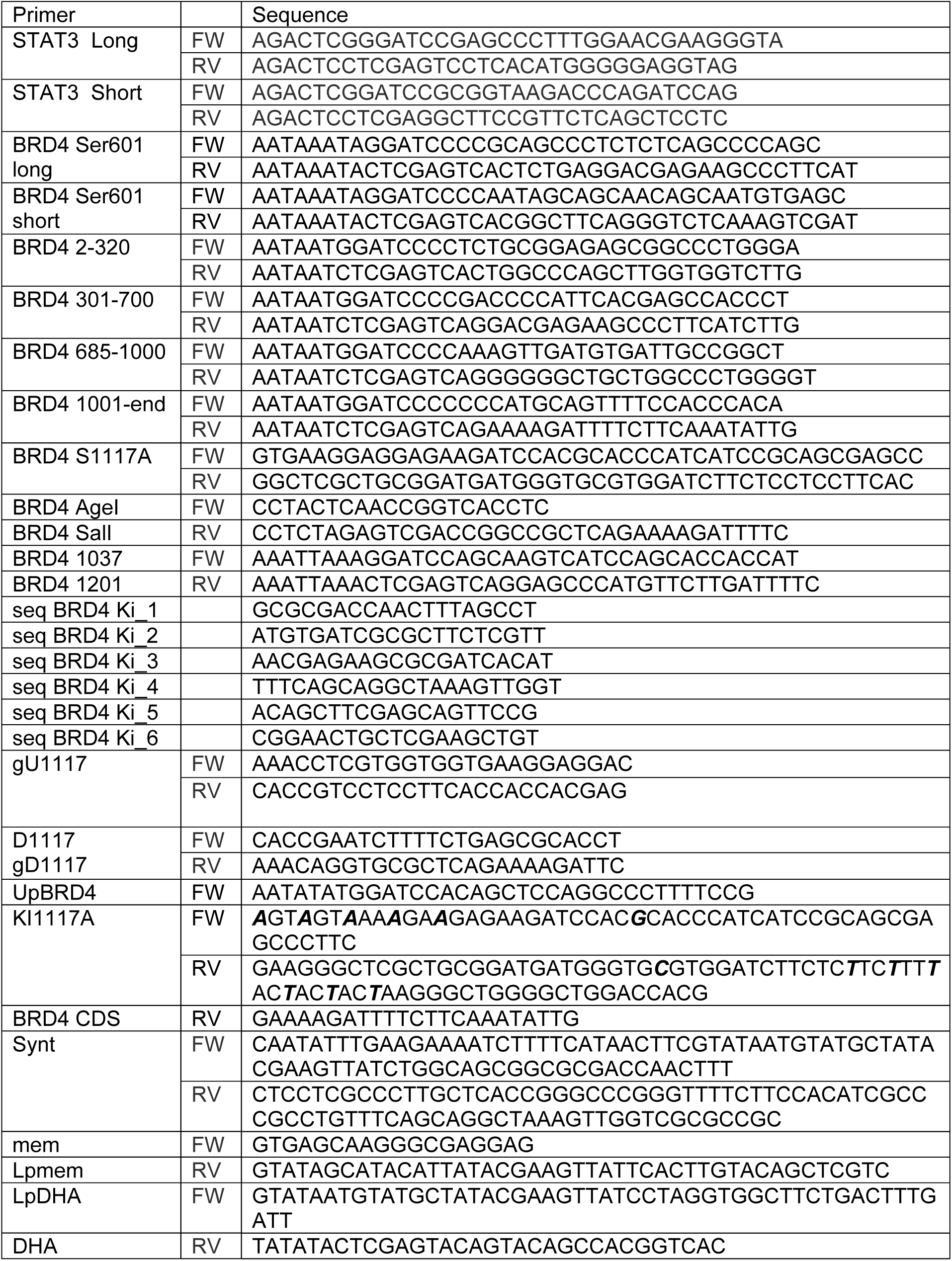
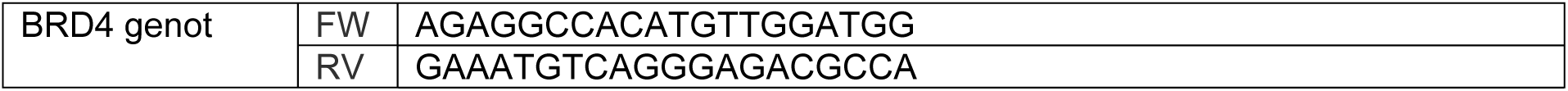
primers used to generate GST fusion proteins, the mutant BRD4 S1117A expression vector and the constructs for gene edition.

**Supplementary Table S3:**
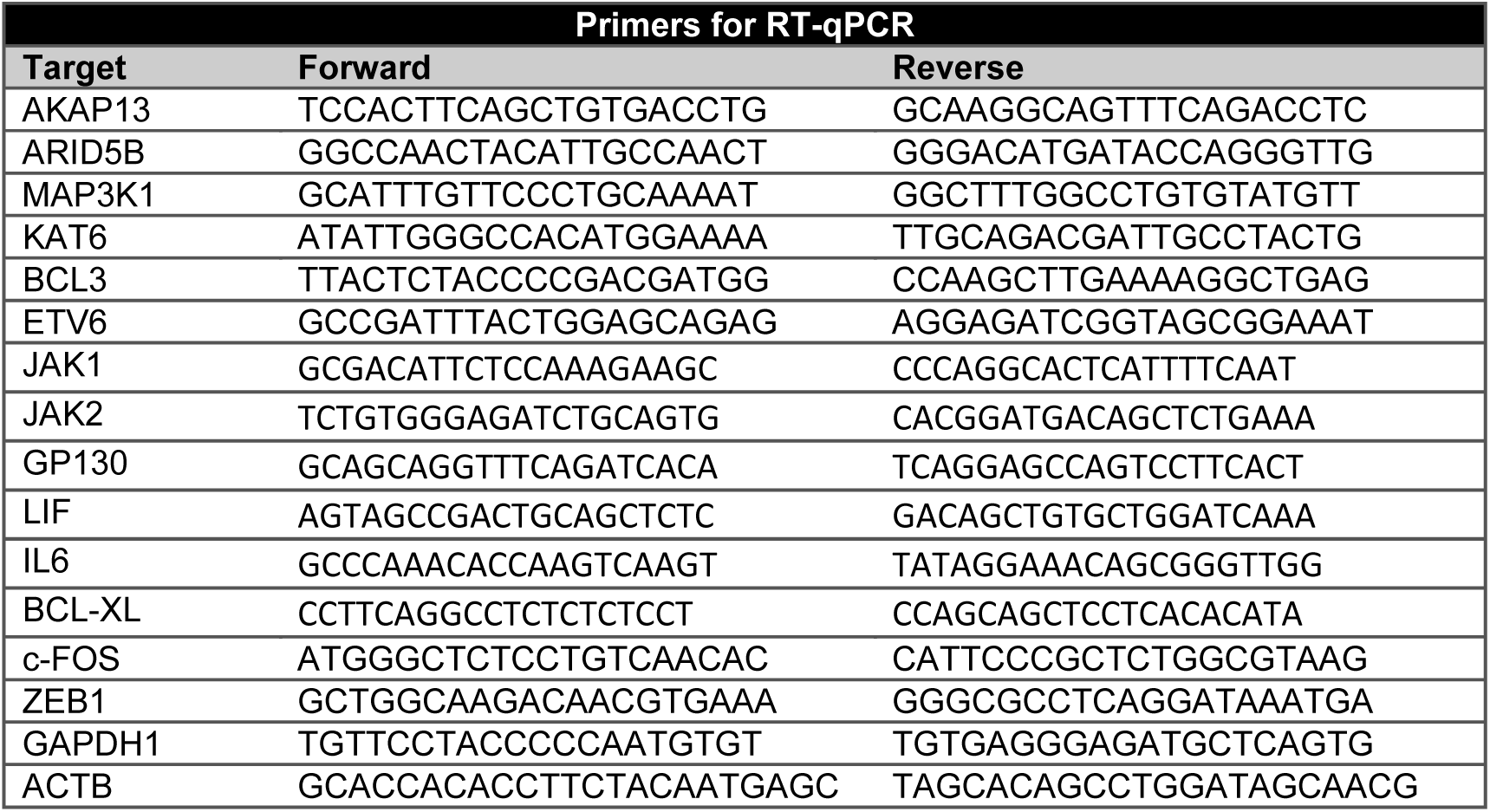
primers used for RT-qPCR analyses.

**Supplementary Table S4:** antibodies used for WB and IHC analyses.

| Antibody | Specie | Dilution | Company | Reference |
| --- | --- | --- | --- | --- |
| BRD4 | Rabbit | 1:1000 | Abcam | ab128874 |
| IKKα | Rabbit | 1:1000 | Cell signaling | #2682s |
| p-IKKα(16A6) | Rabbit | 1:1000 | Cell signaling | #2697 |
| H4 | Rabbit | 1:1000 | Abcam | ab-10158 |
| H4k5Ac | Rabbit | 1:1000 | Active Motif | 39169 |
| Tubulin | Mouse | 1:10000 | Sigma | T6074 |
| Lamin B | Goat | 1:1000 | Santa Cruz | sc-6216 |
| STAT3 | Rabbit | 1:1000 | Cell signaling | #12640s |
| p-STAT3(S727) | Mouse | 1:1000 | Cell signaling | sc-71792 |
| p-STAT3(Y705) | Rabbit | 1:1000 | Cell signaling | #9145s |
| cCAS3 | Rabbit | 1:1000 | Cell signaling | #9661s |
| PARP1 | Mouse | 1:1000 | From Dr. Yélamos (IMIM) |  |
| p-ERK1/2 | Rabbit | 1:1000 | Cell signaling | #9102s |
| ERK1/2 | Mouse | 1:1000 | Milipore | 13-6200 |
| BRD4 ChIP | Rabbit | 2μg/IP | Diagenode | C15410337 |
| Ki67 (MM1) | Mouse | 1:1000 | Leica Biosystems | RRID:AB_442101 |
| LIF (blocking Ab) | Rabbit | 1:200 | Joan Seoane's Lab |  |

## REFERENCES

1. Amado, R. G. et al. Wild-type KRAS is required for panitumumab efficacy in patients with metastatic colorectal cancer. J. Clin. Oncol. 26, 1626–34 (2008).

2. Di Nicolantonio, F. et al. Wild-Type BRAF Is Required for Response to Panitumumab or Cetuximab in Metastatic Colorectal Cancer. J. Clin. Oncol. 26, 5705–5712 (2008).

3. Prescott, J. & Cook, S. Targeting IKKβ in Cancer: Challenges and Opportunities for the Therapeutic Utilisation of IKKβ Inhibitors. Cells 7, 115 (2018).

4. Colomer, C., Marruecos, L., Vert, A., Bigas, A. & Espinosa, L. NF-κB Members Left Home: NF-κB-Independent Roles in Cancer. Biomedicines 5, 26 (2017).

5. Colomer, C. et al. IKKα Kinase Regulates the DNA Damage Response and Drives Chemo-resistance in Cancer. Mol. Cell 75, 669–682.e5 (2019).

6. Dey, A., Chitsaz, F., Abbasi, A., Misteli, T. & Ozato, K. The double bromodomain protein Brd4 binds to acetylated chromatin during interphase and mitosis. Proc. Natl. Acad. Sci. 100, 8758–8763 (2003).

7. Bisgrove, D. A., Mahmoudi, T., Henklein, P. & Verdin, E. Conserved P-TEFb-interacting domain of BRD4 inhibits HIV transcription. Proc. Natl. Acad. Sci. U. S. A. 104, 13690–5 (2007).

8. Devaiah, B. N. et al. BRD4 is a histone acetyltransferase that evicts nucleosomes from chromatin. Nat. Struct. Mol. Biol. 23, 540–548 (2016).

9. Li, X. et al. BRD4 Promotes DNA Repair and Mediates the Formation of TMPRSS2-ERG Gene Rearrangements in Prostate Cancer. Cell Rep. 22, 796–808 (2018).

10. Ni, M. et al. BRD4 inhibition sensitizes cervical cancer to radiotherapy by attenuating DNA repair. Oncogene 40, 2711–2724 (2021).

11. Wakita, M. et al. A BET family protein degrader provokes senolysis by targeting NHEJ and autophagy in senescent cells. Nat. Commun. 11, 1935 (2020).

12. Sun, C. et al. BRD4 Inhibition Is Synthetic Lethal with PARP Inhibitors through the Induction of Homologous Recombination Deficiency. Cancer Cell 33, 401–416.e8 (2018).

13. Zhang, J. et al. BRD4 facilitates replication stress-induced DNA damage response. Oncogene 37, 3763–3777 (2018).

14. Floyd, S. R. et al. The bromodomain protein Brd4 insulates chromatin from DNA damage signalling. Nature 498, 246–50 (2013).

15. Donati, B., Lorenzini, E. & Ciarrocchi, A. BRD4 and Cancer: going beyond transcriptional regulation. Mol. Cancer 17, 164 (2018).

16. Ray, S. et al. Inducible STAT3 NH2 terminal mono-ubiquitination promotes BRD4 complex formation to regulate apoptosis. Cell. Signal. 26, 1445–1455 (2014).

17. Shorstova, T., Foulkes, W. D. & Witcher, M. Achieving clinical success with BET inhibitors as anti-cancer agents. Br. J. Cancer 124, 1478–1490 (2021).

18. Yu, H., Lee, H., Herrmann, A., Buettner, R. & Jove, R. Revisiting STAT3 signalling in cancer: new and unexpected biological functions. Nat. Rev. Cancer 14, 736–746 (2014).

19. Bowman, T. et al. Stat3-mediated Myc expression is required for Src transformation and PDGF-induced mitogenesis. Proc. Natl. Acad. Sci. 98, 7319–7324 (2001).

20. Leslie, K. et al. Cyclin D1 Is Transcriptionally Regulated by and Required for Transformation by Activated Signal Transducer and Activator of Transcription 3. Cancer Res. 66, 2544–2552 (2006).

21. Yu, H. et al. LIF negatively regulates tumour-suppressor p53 through Stat3/ID1/MDM2 in colorectal cancers. Nat. Commun. 5, 5218 (2014).

22. Jones, V. S. et al. Cytokines in cancer drug resistance: Cues to new therapeutic strategies. Biochim. Biophys. Acta 1865, 255–65 (2016).

23. Hu, X., Li, J., Fu, M., Zhao, X. & Wang, W. The JAK/STAT signaling pathway: from bench to clinic. Signal Transduct. Target. Ther. 6, 402 (2021).

24. Darnell, J. E., Kerr, lan M. & Stark, G. R. Jak-STAT Pathways and Transcriptional Activation in Response to IFNs and Other Extracellular Signaling Proteins. Science (80-. ). 264, 1415–1421 (1994).

25. Liu, Y.-N. et al. Leukemia Inhibitory Factor Promotes Castration-resistant Prostate Cancer and Neuroendocrine Differentiation by Activated ZBTB46. Clin. Cancer Res. 25, 4128–4140 (2019).

26. McLean, K. et al. Leukemia inhibitory factor functions in parallel with interleukin-6 to promote ovarian cancer growth. Oncogene 38, 1576–1584 (2019).

27. Peñuelas, S. et al. TGF-β Increases Glioma-Initiating Cell Self-Renewal through the Induction of LIF in Human Glioblastoma. Cancer Cell 15, 315–327 (2009).

28. Pascual-García, M. et al. LIF regulates CXCL9 in tumor-associated macrophages and prevents CD8+ T cell tumor-infiltration impairing anti-PD1 therapy. Nat. Commun. 10, 2416 (2019).

29. Hallett, R. M. et al. Therapeutic Targeting of LIF Overcomes Macrophage-mediated Immunosuppression of the Local Tumor Microenvironment. Clin. Cancer Res. 29, 791–804 (2023).

30. Jang, M. K. et al. The Bromodomain Protein Brd4 Is a Positive Regulatory Component of P-TEFb and Stimulates RNA Polymerase II-Dependent Transcription. Mol. Cell 19, 523–534 (2005).

31. Ai, N. et al. Signal-induced Brd4 release from chromatin is essential for its role transition from chromatin targeting to transcriptional regulation. Nucleic Acids Res. 39, 9592–9604 (2011).

32. Barrows, J. K. et al. BRD4 promotes resection and homology-directed repair of DNA double-strand breaks. Nat. Commun. 13, 3016 (2022).

33. Colomer, C. et al. IKKα is required in the intestinal epithelial cells for tumour stemness. Br. J. Cancer 118, 839–846 (2018).

34. Carpenter, R. & Lo, H.-W. STAT3 Target Genes Relevant to Human Cancers. Cancers (Basel). 6, 897–925 (2014).

35. Robinson, R. L. et al. Comparative STAT3-Regulated Gene Expression Profile in Renal Cell Carcinoma Subtypes. Front. Oncol. 9, 72 (2019).

36. Brocke-Heidrich, K. et al. BCL3 is induced by IL-6 via Stat3 binding to intronic enhancer HS4 and represses its own transcription. Oncogene (2006) doi:10.1038/sj.onc.1209711.

37. Chang, T. P. & Vancurova, I. Bcl3 regulates pro-survival and pro-inflammatory gene expression in cutaneous T-cell lymphoma. Biochim. Biophys. Acta - Mol. Cell Res. (2014) doi:10.1016/j.bbamcr.2014.07.012.

38. Urban, B. C. et al. BCL-3 expression promotes colorectal tumorigenesis through activation of AKT signalling. Gut (2016) doi:10.1136/gutjnl-2014-308270.

39. Zou, Y. et al. The proto-oncogene Bcl3 induces immune checkpoint PD-L1 expression, mediating proliferation of ovarian cancer cells. J. Biol. Chem. 293, 15483–15496 (2018).

40. Margalef, P. et al. BRAF-induced tumorigenesis is IKKα-dependent but NF-κB–independent. Sci. Signal. 8, ra38–ra38 (2015).

41. Verhoeven, Y. et al. The potential and controversy of targeting STAT family members in cancer. Semin. Cancer Biol. 60, 41–56 (2020).

42. Delmore, J. E. et al. BET Bromodomain Inhibition as a Therapeutic Strategy to Target c-Myc. Cell 146, 904–917 (2011).

43. Marisa, L. et al. Gene Expression Classification of Colon Cancer into Molecular Subtypes: Characterization, Validation, and Prognostic Value. PLoS Med. (2013) doi:10.1371/journal.pmed.1001453.

44. Guinney, J. et al. The consensus molecular subtypes of colorectal cancer. Nat. Med. 21, 1350–1356 (2015).

45. Llosa, N. J. et al. The vigorous immune microenvironment of microsatellite instable colon cancer is balanced by multiple counter-inhibitory checkpoints. Cancer Discov. 5, 43–51 (2015).

46. Itzen, F., Greifenberg, A. K., Bösken, C. A. & Geyer, M. Brd4 activates P-TEFb for RNA polymerase II CTD phosphorylation. Nucleic Acids Res. 42, 7577–7590 (2014).

47. Oliveira, A. F., Bretes, L. & Furtado, I. Review of PD-1/PD-L1 Inhibitors in Metastatic dMMR/MSI-H Colorectal Cancer. Front. Oncol. 9, 396 (2019).

48. Hahn, Y.-I. et al. STAT3 Stabilizes IKKα Protein through Direct Interaction in Transformed and Cancerous Human Breast Epithelial Cells. Cancers (Basel). 13, 82 (2020).

49. Christianson, J., Oxford, J. T. & Jorcyk, C. L. Emerging Perspectives on Leukemia Inhibitory Factor and its Receptor in Cancer. Front. Oncol. 11, 693724 (2021).

50. Colomer, C., Pecharroman, I., Bigas, A. & Espinosa, L. Targeting IKKα kinase to prevent tumor progression and therapy resistance. Cancer Drug Resist. (2020) doi:10.20517/cdr.2019.104.

51. Sato, T. et al. Long-term Expansion of Epithelial Organoids From Human Colon, Adenoma, Adenocarcinoma, and Barrett’s Epithelium. Gastroenterology 141, 1762–1772 (2011).

52. Espinosa, L., Inglés-Esteve, J., Aguilera, C. & Bigas, A. Phosphorylation by Glycogen Synthase Kinase-3β Down-regulates Notch Activity, a Link for Notch and Wnt Pathways. J. Biol. Chem. 278, 32227–32235 (2003).

53. Krueger, F. Trim Galore. (2012).

54. Langmead, B. & Salzberg, S. L. Fast gapped-read alignment with Bowtie 2. Nat. Methods 9, 357–9 (2012).

55. Zhang, Y. et al. Model-based Analysis of ChIP-Seq (MACS). Genome Biol. 9, R137 (2008).

56. Sloan, C. A. et al. ENCODE data at the ENCODE portal. Nucleic Acids Res. 44, D726–D732 (2016).

57. Yu, G., Wang, L.-G. & He, Q.-Y. ChIPseeker: an R/Bioconductor package for ChIP peak annotation, comparison and visualization. Bioinformatics 31, 2382–2383 (2015).

58. Cavalcante, R. G. & Sartor, M. A. Annotatr: Genomic regions in context. Bioinformatics 33, 2381–2383 (2017).

59. Kuleshov, M. V et al. Enrichr: a comprehensive gene set enrichment analysis web server 2016 update. Nucleic Acids Res. 44, W90–7 (2016).

60. Ramírez, F. et al. deepTools2: a next generation web server for deep-sequencing data analysis. Nucleic Acids Res. 44, W160–W165 (2016).

61. Love, M. I., Huber, W. & Anders, S. Moderated estimation of fold change and dispersion for RNA-seq data with DESeq2. Genome Biol. 15, 550 (2014).

62. Colaprico, A. et al. TCGAbiolinks: an R/Bioconductor package for integrative analysis of TCGA data. Nucleic Acids Res. 44, e71–e71 (2016).

63. Sanjana, N. E., Shalem, O. & Zhang, F. Improved vectors and genome-wide libraries for CRISPR screening. Nat. Methods 11, 783–784 (2014).

64. Espinosa, L., Inglés-Esteve, J., Robert-Moreno, A. & Bigas, A. IκBα and p65 Regulate the Cytoplasmic Shuttling of Nuclear Corepressors: Cross-talk between Notch and NFκB Pathways. Mol. Biol. Cell 14, 491–502 (2003).

